# Maternal vaccination timing shapes the composition and maturation of antibody clonotypes transferred to the newborn

**DOI:** 10.64898/2026.07.13.738171

**Authors:** Aisha Yesbolatova, Lowrey Peyton, Nicholas C. Curtis, Tae Kim, Jinyoung Yoon, Hunter J. Melton, Noor M. Taher, Tae-Geun Yu, Brian H. Kim, Dhvanir N. Kansara, Matthew R. Stoner, Ellison Ober, Emma Varella, Pieter Pannus, Larissa DeBrabandere, Kirsten Maertens, Kenneth B. Hoehn, Mark Connors, Arnaud Marchant, Margaret E. Ackerman, Jiwon Lee

## Abstract

Newborns rely on maternally transferred antibodies for immune protection, acquired across the placenta and through breast milk. Despite this importance, how closely the inherited antibody repertoire resembles the mother’s has not been examined at the clonotypic level. We combined BCR-Seq with proteomic Ig-Seq to track SARS-CoV-2-specific antibody clonotypes across maternal blood, cord blood, and breast milk from six mRNA-immunized pregnant individuals. Vaccination earlier in gestation generated more diverse peak IgG repertoires but greater contraction before delivery, yielding fewer transferred clonotypes. Vaccination later in gestation produced more restricted peak repertoires, but more clonotypes persisted to delivery and transferred to cord. Infant cord was enriched for persistent, highly somatically mutated, and intraclonally diverse IgG clonotypes, consistent with preferential transfer of affinity-matured antibodies. In contrast, breast milk IgA repertoires were largely distinct from systemic repertoires and underwent substantial remodeling despite stable antigen titers. These findings define molecular determinants of passive immunity shaped by vaccination timing relevant to optimizing maternal immunization.

## Introduction

Newborns are most vulnerable to infectious diseases in the first few months of life, during which maternally derived antibodies are a principal determinant of infant immunity^1^. Maternal antibody transfer begins *in utero* via transplacental IgG transport and continues postnatally through breastfeeding, creating complementary prenatal and postpartum routes of passive protection before infants establish robust endogenous humoral immunity^2^.

Accordingly, maternal vaccination during pregnancy is a critical public health strategy for protecting both pregnant individuals and infants. Over the course of the SARS-CoV-2 pandemic, mRNA vaccination during pregnancy was associated with reduced maternal infection risk^3–6^ and with reduced risks of SARS-CoV-2 infection and COVID-19–related hospitalization in infants^7–9^, with infant protection correlating with cord blood IgG levels^10^. Spike (S)-specific antibodies are also present in human milk after infection or vaccination and persist for up to one year^11–14^, providing a distinct, ongoing source of antibody to the infant mucosa during breastfeeding.

Transplacental IgG transfer is mediated primarily by the neonatal Fc receptor (FcRn) expressed on syncytiotrophoblasts, which binds IgG at the acidic pH of the endosome and releases it at neutral pH on the fetal side^15^. IgG subclasses are not transferred equally, and transfer efficiency varies with gestational age and other factors^16,17^, with subclass transfer hierarchy and factors that modify transfer efficiency documented in recent reviews^18,19^. Beyond subclass, systems serology and modeling studies suggest that placental transfer can be functionally selective, with Fc receptor binding and Fc glycan patterns associating with differential enrichment of antibody features in cord blood^20^, although whether Fc glycan composition directly governs transfer efficiency remains contested^15^. FcRn expression in the placenta increases with placental maturation in the second and third trimesters^21^, and infant IgG levels at birth often exceed matched maternal levels, reflecting accumulation as placental transfer capacity increases. Maternal antibody abundance in late gestation is therefore a central determinant of the amount of antigen-specific IgG available for transfer^22–24^.

Although the quantitative determinants of placental antibody transfer are increasingly well characterized, they are limited to understanding how much antibody reaches the newborn rather than which antibodies transfer. Prior work has shown that transfer is selective at the level of Fc features and IgG subclasses, yet whether this selectivity extends to the antigen-specific clonotypes remains unknown. Consequently, the sequence-level features that distinguish antibody lineages inherited by the newborn have not been defined. Standard serology quantifies maternal and cord antibody levels but does not resolve which clonotypes are transferred. Because maternal vaccination can occur across a wide range of gestational ages, it is also unknown whether vaccination timing influences the molecular composition, and not only the quantity, of the antibodies that reach the newborn.

The breast milk antibody compartment is even less well defined at the molecular level. The antibody response in human milk varies with the type of infection or vaccine, where immunity was induced, and maternal factors such as geographic location and body mass index^25^. Secreted antibody in milk is dominated by secretory IgA produced locally by plasma cells that home to the mammary gland.

Proteomics-based profiling indicates that milk IgA and serum IgG1 repertoires are largely distinct^26,27^, and likely arise from compartmentalized antibody production^28^. Yet the extent to which vaccine-elicited systemic responses give rise to clonotypes in the milk compartment, and how the clonotypic composition of the secreted antibody repertoire in milk changes over the course of lactation, remain unknown. Whether the antibodies an infant receives through breast milk are drawn from the systemic maternal response or generated independently at the mucosa can be resolved only by comparing the milk, serum, and cord antibody repertoires at the clonotypic level within the same mother-infant dyads.

We report here the clonotype-level deconvolution of vaccine-elicited maternal antibody repertoires across maternal serum, umbilical cord blood, and breast milk following two doses of BNT162b2. Our results reveal that earlier vaccination generated a more diverse peak maternal IgG repertoire, but the longer interval before delivery resulted in greater clonotype contraction, so fewer lineages persisted to the late-gestation transfer window. Later vaccination produced a more restricted peak repertoire but one that was more fully retained at delivery, yielding greater clonotype-level representation in cord blood. Across donors, clonotypes enriched in cord blood showed higher somatic hypermutation and greater intraclonal diversity than those absent from cord blood. Breast milk IgA repertoires were largely distinct from the systemic serological repertoires and were remodeled over the course of lactation. Together, these findings define how vaccination timing and compartment-specific immunoglobulin secretion shape the molecular composition of passively transferred neonatal immunity.

## Results

### Study design

To define the molecular composition of vaccine-elicited maternal antibody repertoires during pregnancy and the sequence-level features associated with transfer to the newborn, we studied six infection-naïve pregnant donors who received two doses of BNT162b2 during pregnancy and delivered at full term (**Fig. 1a,b** and **Supplementary Table 1**). Maternal peripheral blood was collected immediately before the second vaccine dose (baseline; M_D0), 28 days after the second dose (peak response; M_D28), and at delivery (M_Deliv), with paired cord blood obtained at delivery. Peripheral blood mononuclear cells (PBMCs) were collected seven days after the second dose to capture the peak post-boost plasmablast response, and breast milk was collected at 4 and 12 weeks postpartum.

**Figure 1.**
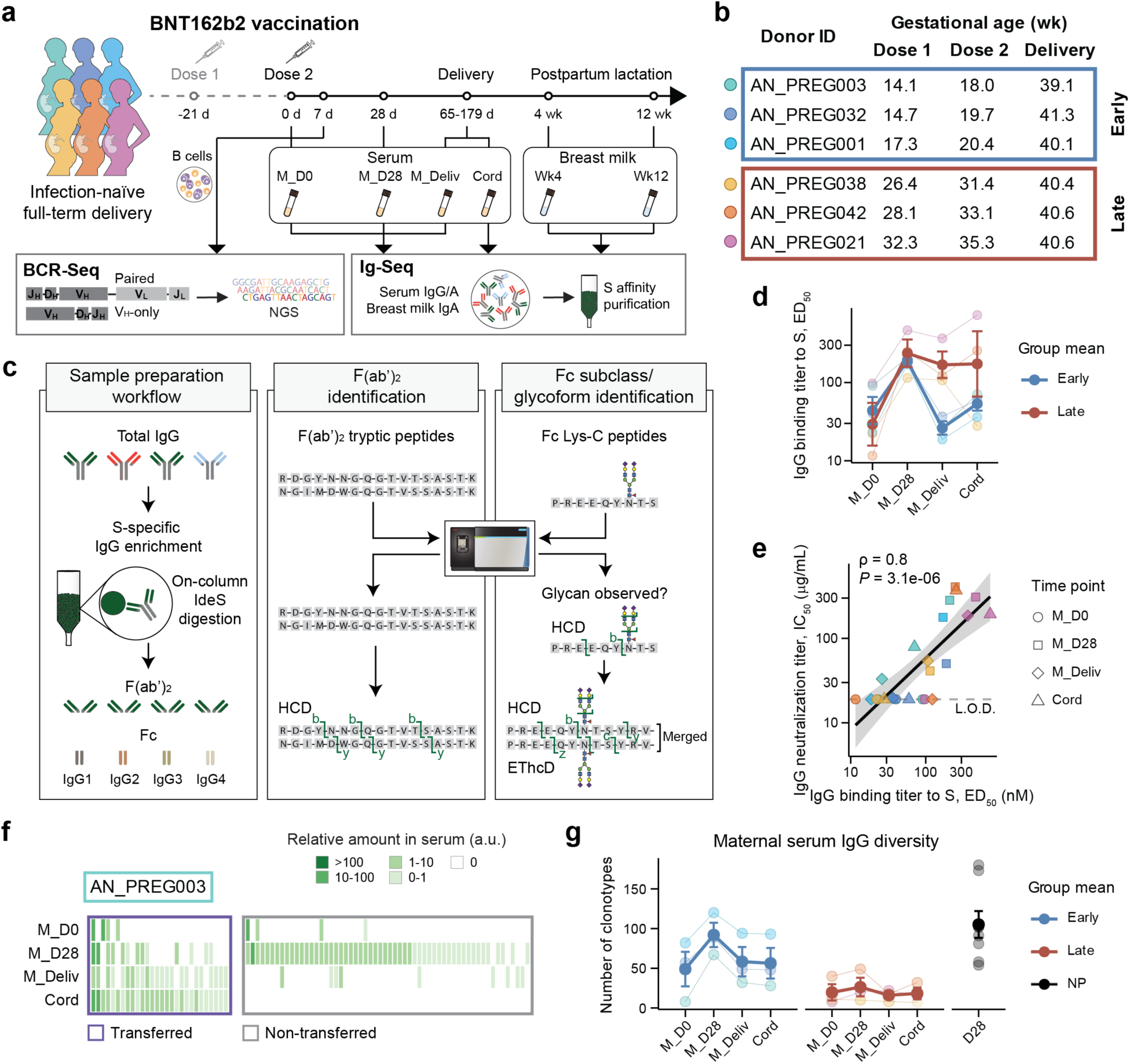
Longitudinal profiling of maternal antibody clonotypes elicited by BNT162b2 vaccination during pregnancy. **a,** Study design and analytical workflow integrating Ig-Seq and BCR-Seq to track vaccine-elicited maternal antibody clonotypes across pregnancy, delivery, and lactation. **b,** Gestational ages at dose 1, dose 2, and delivery for each donor. Donors are grouped into early and late vaccination groups by gestational age at dose 1. **c,** LC-MS/MS workflow for molecular deconvolution of S-reactive serum IgG repertoire clonotypes and for Fc subclass and glycoform profiling. S-reactive IgG was affinity-purified on S-conjugated resin and cleaved by IdeS into F(ab’)_2_ and Fc fragments for clonotype identification or subclass and glycoform assignment, respectively. **d,** S-reactive IgG binding responses as measured by ELISA (ED_50_) values of binding to S in maternal serum at the indicated time points and in paired cord blood. Thin lines represent individual donors, and thick lines represent group means, with error bars indicating standard error of the mean (s.e.m.). **e,** Association between IgG binding titer to S and neutralization across matched samples from all donors and time points. Symbols denote sampling time points. The regression line is shown with 95% confidence interval (gray shading). The dashed line indicates the assay limit of detection (L.O.D.). **f,** Representative heatmap from donor AN_PREG003 showing the relative serum abundance of IgG clonotypes comprising the S-specific serological repertoire over time. Each column represents an individual IgG clonotype, with color indicating its relative amount in serum (arbitrary units), calculated by scaling the anti-S serum IgG binding titer by the relative abundance of each clonotype determined by proteomic analysis. Clonotypes were classified as transferred (detected in cord) or non-transferred (absent in cord). **g,** S-specific IgG clonotype diversity (number of detected clonotypes) in maternal serum over time for early and late vaccination groups, with a non-pregnant (NP) BNT162b2-vaccinated cohort of 8 donors shown at the post-boost time point (D28). Thin lines represent individual donors, and thick lines represent group means, with error bars indicating s.e.m.

To resolve clonotype-level changes in the maternal antibody response, we integrated BCR-Seq and Ig-Seq analyses^29–32^. We recovered both single-chain heavy-chain variable region (V_H_) sequences and natively paired V_H_ and light-chain variable region (V_H_:V_L_) sequences from PBMCs (**Supplementary Table 2**), which were used to construct donor-specific search databases for proteomic assignment of serum IgG clonotypes. Spike (S)-specific serum IgG was affinity-purified, processed for high-resolution liquid chromatography tandem mass spectrometry (LC-MS/MS), and assigned at the clonotype level by matching CDRH3-containing peptides to the corresponding donor-specific search databases (**Supplementary Table 3**). In parallel, Fc domains of the same S-specific IgG fractions were separately analyzed to define subclass and Fc glycoform compositions (**Fig. 1c**). Across all donors and samples, we injected over 300 replicates for LC-MS/MS analysis, taking almost 1,000 hours of injection time alone.

### Longitudinal dynamics of maternal vaccine-elicited serum IgG repertoires

Pregnant donors were stratified by gestational age at vaccination into early and late immunization groups (mean gestational age at dose 1, 15.4 and 28.9 weeks, respectively; **Fig. 1b**). The two groups had similar maternal S-reactive IgG binding titers (measured as 50% Effective Dilution, ED_50_) at M_D0 and M_D28, indicating comparable systemic responses shortly after vaccination (**Fig. 1d**). By delivery, maternal serum titers in the early group had declined from the M_D28 peak, consistent with waning over the longer interval between vaccination and parturition. Despite this decline, cord titers in early-vaccinated dyads remained detectable and exceeded matched maternal titers at delivery in most dyads, consistent with accumulation of vaccine-elicited IgG over the prolonged period of maternal seropositivity. In contrast, cord titers in the late immunization group more closely paralleled maternal titers at delivery. Across maternal serum and cord samples, S-reactive IgG binding titers correlated strongly with neutralization activity (**Fig. 1e** and **Supplementary Fig. 1**).

For Ig-Seq analyses, maternal serum IgG clonotypes were classified as transferred clonotypes (detected in cord), and non-transferred clonotypes (not detected in cord) (**Fig. 1f** and **Supplementary Fig. 2**). The maternal S-specific serum IgG repertoire comprised between 12 and 145 clonotypes across donors, whereas paired cord repertoires contained between 7 and 93 clonotypes. Early vaccination was associated with greater maternal clonotype diversity than late vaccination, most prominently at M_D28 (mean 92 vs. 26 clonotypes). Diversity in the early immunization group was comparable to that in a cohort of non-pregnant mRNA-vaccinated recipients (**Supplementary Table 4**) at the equivalent post-boost time point (**Fig. 1g**). Vaccination timing was therefore associated with both the magnitude of the maternal response and the clonal diversity available for placental transfer.

### Sequence-level features associated with selective placental transfer of maternal IgG clonotypes

To identify features associated with selective placental transfer, we examined the relationship between maternal and cord S-reactive IgG titers. In this cohort, cord blood titers were strongly correlated with peak maternal titers at M_D28 (Spearman’s correlation ⍴ = 1, adjusted *P* = 0.013) but not with maternal titers at delivery (Spearman’s correlation ρ = 0.6, adjusted *P* = 0.338) (**Fig. 2a** and **Supplementary Fig. 3**). This pattern was consistent for neutralization activity, with the peak maternal response strongly correlated with cord neutralization (Spearman’s correlation ρ = 0.94, adjusted *P* = 0.018), but not maternal neutralization at delivery (Spearman’s correlation ρ = 0.03, adjusted *P* = 0.954) (**Supplementary Fig. 3**). Given the small number of dyads, these correlations should be interpreted with caution, but together they indicate that the magnitude of the peak post-boost response was more closely associated with cord S-specific IgG titer than the residual maternal antibody response at parturition.

**Figure 2.**
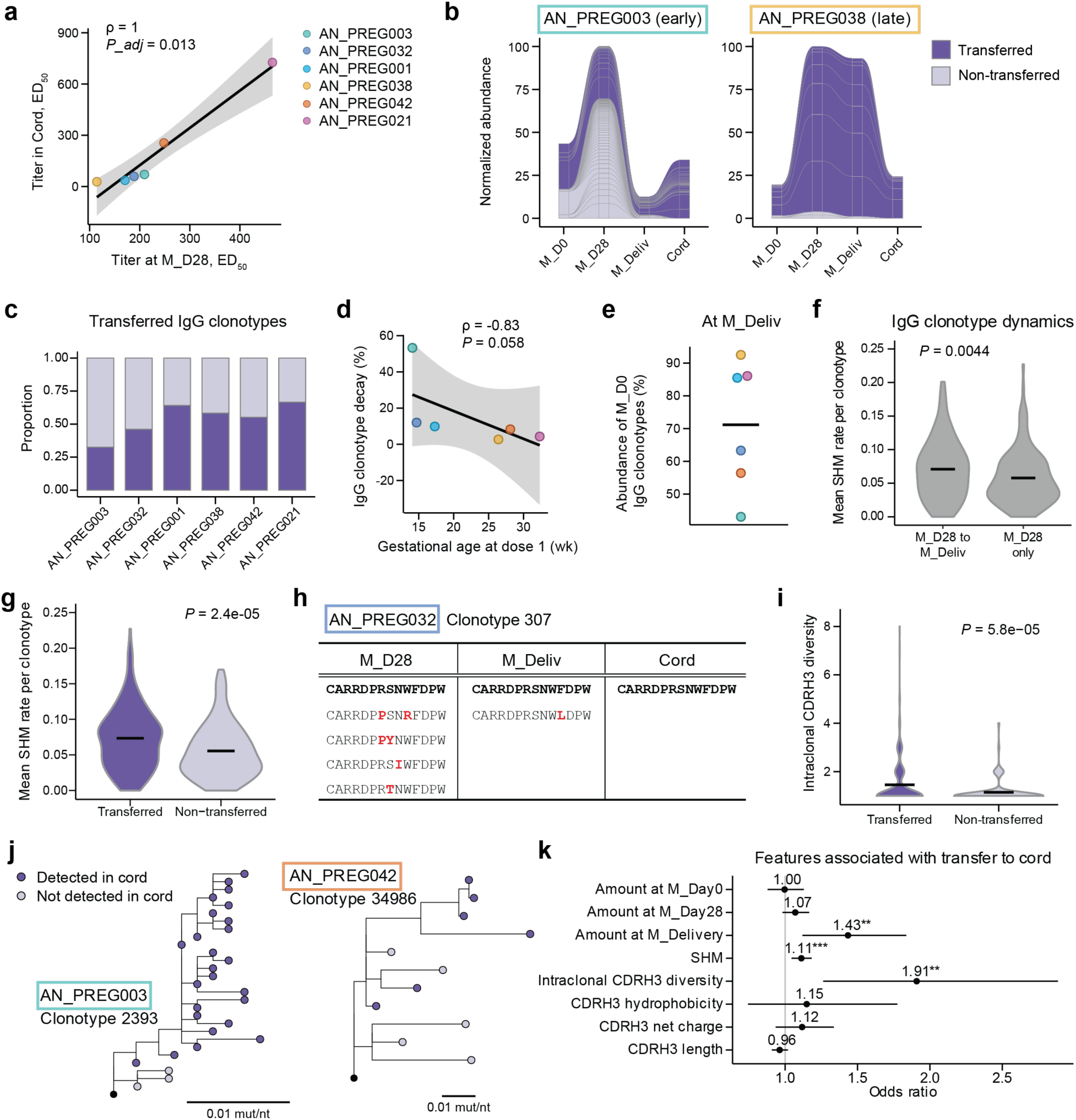
Clonotype features associated with placental transfer of maternal IgG. **a,** Correlation between S-specific IgG binding titer in maternal serum at M_D28 and paired cord at delivery. The regression line is shown with 95% confidence interval (gray shading). **b,** Representative alluvial plots from an early-vaccinated donor (AN_PREG003) and a late-vaccinated donor (AN_PREG038) showing the transferred and non-transferred IgG clonotypes over time, normalized to M_D28. **c,** Proportion of transferred and non-transferred IgG clonotypes in each donor. **d,** Relationship between gestational age at dose 1 and clonotypic decay, defined as the decline in cumulative relative abundance of transferred IgG clonotypes between M_D28 and M_Deliv. The regression line is shown with 95% confidence interval (gray shading). **e,** Fraction of S-reactive IgG clonotypes present at M_D0 that persisted to M_Deliv. **f,** Mean V_H_ SHM for IgG clonotypes that persisted from M_D28 to M_Deliv (persistent) and those detected only at M_D28 (transient). **g,** Mean V_H_ SHM for transferred vs. non-transferred IgG clonotypes. **h,** CDRH3 peptide sequences detected across time points for a representative transferred clonotype (AN_PREG032 clonotype 307), illustrating intraclonal variants. Residues that differ from the most abundant CDRH3 variant are shown in red. **i,** Intraclonal CDRH3 diversity (number of unique CDRH3 peptides per clonotype) for transferred and non-transferred IgG clonotypes. **j,** Representative phylogenetic trees of two B cell clonal lineages. Purple circles indicate lineage members whose CDRH3 matches a clonotype detected in cord, and gray circles indicate members not detected in cord. UCA, unmutated common ancestor. Branches are scaled to mutations per nucleotide site. **k,** Odds ratios from a GLMM of clonotype-level predictors of transfer to cord. Vertical line represents null value. Asterisks indicate significance (*p<0.05, **p<0.01, ***p<0.001). Throughout figure, crossbars indicate mean.

Clonotype-level deconvolution of the serological IgG repertoire revealed markedly distinct transfer dynamics between vaccination timing groups. Early-vaccinated donors showed a large expansion of S-specific IgG clonotypes at M_D28 followed by pronounced contraction before delivery, such that many clonotypes present at peak were not detected in cord (**Fig. 2b** and **Supplementary Fig. 4**). Late-vaccinated donors retained a larger fraction of M_D28 clonotypes through delivery, and these persistent clonotypes were more often represented in cord. Accordingly, the early group had a lower proportion of maternal clonotypes detected in cord than the late group (group means 47.6% vs. 60.1%; **Fig. 2c**). Quantitatively, clonotypic decay, defined as the decline in cumulative abundance of transferred clonotypes between M_D28 and M_Deliv, decreased as gestational age at dose 1 increased (Spearman’s correlation ⍴ = -0.83, *P* = 0.058; **Fig. 2d**), consistent with reduced repertoire contraction when vaccination occurred later in pregnancy.

We next examined whether persistence to delivery marked clonotypes more likely to be transferred. On average, 71% of the IgG clonotypes present at M_D0 persisted to M_Deliv (**Fig. 2e**), indicating a major contribution from pre-existing clonotypes recalled by the second dose. IgG clonotypes that persisted from M_D28 to M_Deliv had higher mean V_H_ somatic hypermutation (SHM) levels than clonotypes detected only at M_D28 (7.1% vs. 5.8% respectively, *P* = 0.0044; **Fig. 2f**). Likewise, transferred IgG clonotypes were more mutated than non-transferred clonotypes (7.3% vs. 5.6% respectively, *P* = 2.4e-05; **Fig. 2g**), consistent with preferential representation of more affinity-matured maternal IgG clonotypes among those detected in cord.

Because SHM alone does not capture within-lineage diversification, we quantified intraclonal diversity as the number of unique CDRH3 peptides detected within each serum IgG clonotype across time points, as described previously^33^. A representative transferred clonotype from an early vaccinee (AN_PREG032, clonotype 307) contained multiple intraclonal variants tracked across M_D28, M_Deliv, and cord (**Fig. 2h**). Overall, transferred clonotypes exhibited greater intraclonal CDRH3 diversity than non-transferred clonotypes (1.46 vs. 1.15; **Fig. 2i**). In specific examples, we observed lineage trees showing cord-detected CDRH3 variants occupying more mutated branches within B cell phylogenies (**Fig. 2j; Supplementary Fig. 5**). *IGHV* gene usage and CDRH3 biophysical properties were similar between transferred and non-transferred repertoires (**Supplementary Fig. 6**).

To identify clonotypic features associated with transfer to cord, we modeled transfer to cord with a generalized linear mixed-effects model (GLMM) using maternal serum IgG abundance across time points, CDRH3 biophysical properties, SHM and intraclonal diversity as predictors and donor identity as a random effect. Maternal clonotype abundance at delivery, mean SHM, and intraclonal CDRH3 diversity were each significantly positively associated with transfer, whereas abundance at M_D0 or M_D28 and CDRH3 biophysical properties were not statistically significant: odds ratio 1.43 (1.12-1.84) for clonotype abundance at delivery; 1.11 (1.04-1.18) for SHM; and 1.91 (1.26-1.88) for intraclonal CDRH3 diversity (**Fig. 2k** and **Supplementary Fig. 7**). These data indicate that placental transfer is selective at the clonotype level and preferentially enriches for persistent maternal IgG clonotypes with molecular features consistent with affinity maturation. That abundance at delivery, but not at M_D28, predicted transfer indicates that representation in cord is determined by which clonotypes remain abundant at parturition rather than by the size of the peak response, and that among those persistent clonotypes, transfer is associated with molecular features of affinity maturation.

### S-specific IgG subclass and Fc glycoform features during pregnancy and in transfer

To assess IgG Fc glycosylation dynamics throughout pregnancy and in transplacental transfer, we performed subclass-resolved analysis of *N*-linked glycosylation by LC-MS/MS of Lys-C-cleaved, S-specific IgG Fc glycopeptides (**Fig. 1c**) using longitudinal maternal samples and matched cord samples from four dyads. Fc glycan analysis was restricted to four donors due to sample volume constraints that precluded inclusion of all six dyads. Unsupervised hierarchical clustering of glycoform abundances (**Supplementary Fig. 8a**) revealed that IgG subclass and donor identity were the dominant sources of glycoform/sample variability (PERMANOVA *R*^2^ effect sizes 0.37 and 0.50, respectively) (**Supplementary Fig. 8b**), indicating that inter-donor differences outweigh longitudinal changes within donors. In maternal-cord dyads at delivery, IgG1 was most enriched in cord followed by IgG3 and IgG4, and IgG2 transferred least efficiently (**Supplementary Fig. 8c**).

Longitudinally, fucosylation, sialylation, and bisection of S-specific IgG Fc glycans declined toward delivery, while cord IgG was relatively enriched for sialylated forms (mean increase of 3.6% between M_Deliv and cord) (**Supplementary Fig. 8d**). A linear mixed-effects model incorporating subclass and glycosylation features identified maternal serum abundance at delivery as the strongest significant predictor of cord abundance, estimate 1.47 (1.39-1.55) (**Supplementary Fig. 8e** and **Supplementary Fig. 9**). No individual glycan feature reached significance. Within this cohort, the amount of each S-specific IgG subclass-glycoform present in maternal serum at parturition was therefore a stronger determinant of transfer than its Fc glycan composition.

### Vaccine-elicited breast milk IgA repertoires are compartmentalized from the systemic serological repertoires

S-reactive breast milk IgA titers were generally low and stable from 4 weeks (Wk4) to 12 weeks (Wk12) postpartum, although one donor (AN_PREG032) who had confirmed SARS-CoV-2 infection between Wk4 and Wk12 had a marked increase at Wk12 (**Fig. 3a** and **Supplementary Table 5**). Neutralizing activity was undetectable in most milk samples and was observed only in the post-infection Wk12 sample from AN_PREG032 (**Fig. 3b**).

**Figure 3.**
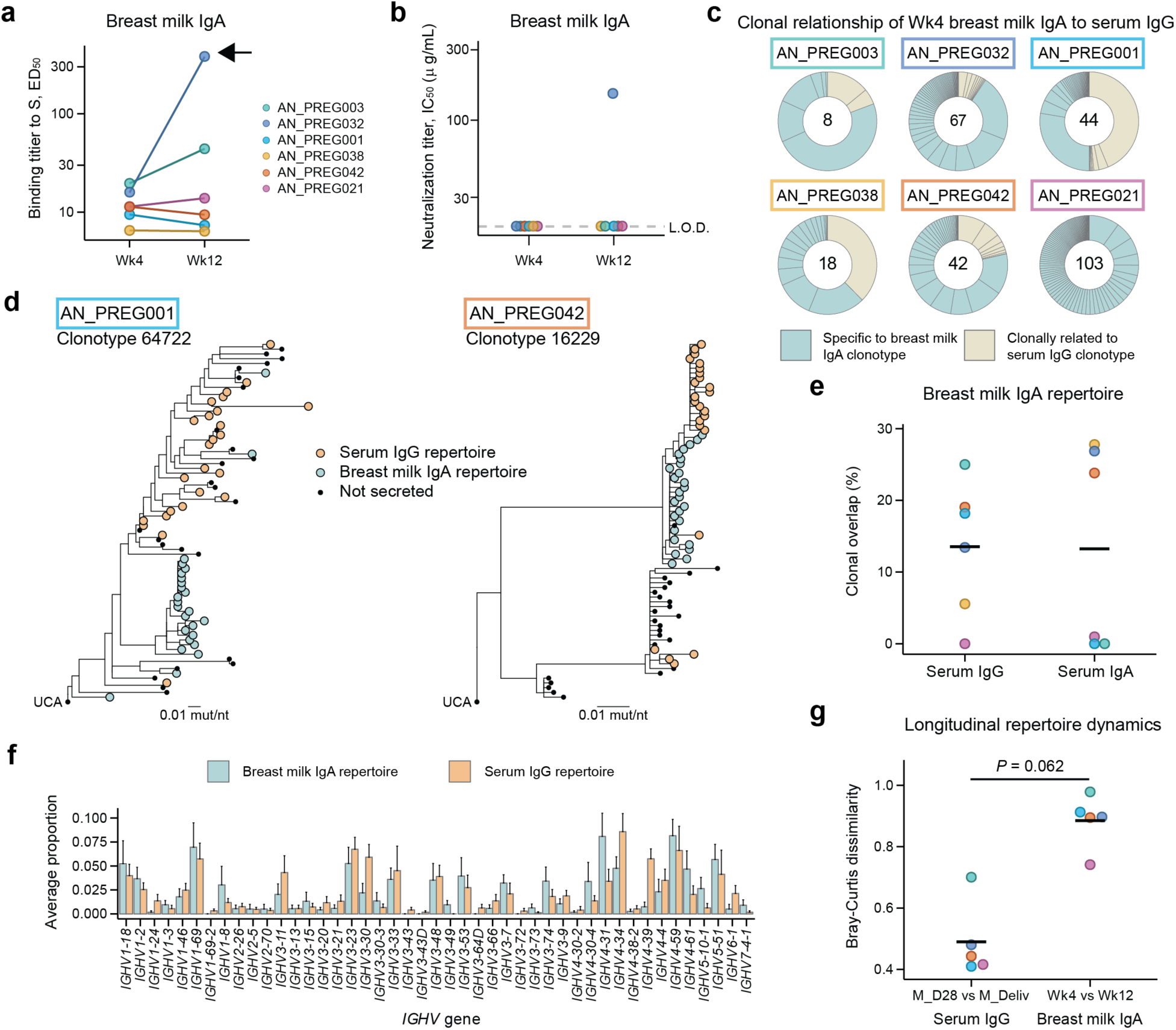
Breast milk IgA repertoires are compartmentalized from systemic maternal antibody repertoires. **a,** S-reactive IgA binding titers in breast milk at Wk4 and Wk12. The arrow indicates donor AN_PREG032, who had a SARS-CoV-2 breakthrough infection between Wk4 and Wk12. **b,** Neutralization activity in breast milk at Wk4 and Wk12. **c,** Donut plots showing donor-specific breast milk IgA repertoires at Wk4. Each segment represents an individual breast milk IgA clonotype, with segment area proportional to relative abundance. Number in the center indicates the total clonotypes detected. **d,** Phylogenetic trees of two B cell clonal lineages from AN_PREG001 and AN_PREG042 that contain members matched to both serum and breast milk repertoires. Tip colors indicate repertoire assignment by CDRH3 match (serum IgG, breast milk IgA, or not secreted). Branch lengths are scaled to mutations per nucleotide site. UCA, unmutated common ancestor. **e,** Frequency of clonal overlap between breast milk IgA clonotypes and maternal serum IgG or serum IgA clonotypes. Crossbar indicates mean. **f,** *IGHV* gene usage frequency in breast milk IgA and serum IgG repertoires. Bars indicate mean proportions across donors; error bars indicate s.e.m. All pairwise comparisons not significant after multiple hypothesis correction. **g,** Longitudinal repertoire dynamics in serum IgG and breast milk IgA compartments, quantified as Bray-Curtis dissimilarity between M_D28 and M_Deliv (serum IgG) or Wk4 and Wk12 (breast milk IgA). Comparison was performed using paired Wilcox test without donor AN_PREG038 due to missing breast milk Wk12 sample. Crossbar indicates mean.

To define the relationship between milk and systemic antibody compartments, we purified IgA from delipidated Wk4 and Wk12 breast milk samples (**Supplementary Fig. 10a**). The purified breast milk IgA was predominantly secretory IgA (sIgA) (**Supplementary Fig. 10b**) and was subjected to S-specific affinity purification. The S-specific breast milk IgA repertoire comprised 8-103 clonotypes per donor, and only a small subset of breast milk IgA clonotypes were clonally related to maternal serum IgG clonotypes (total 28/282 IgA clonotypes overlapped, 9.9%, across all donors; **Fig. 3c**, **Supplementary Fig. 10c**). A few of these rare shared clonotypes nonetheless accounted for a substantial fraction of breast milk IgA abundance. Specific examples of phylogenetic trees of shared clonotypes showed that serum IgG- and breast milk IgA-matched CDRH3 sequences occupied largely distinct branches descending from common ancestors (**Fig. 3d**), consistent with early divergence followed by compartment-specific diversification. Clonal overlap frequency was comparably low when breast milk IgA clonotypes were compared with either serum IgG or serum IgA clonotypes (**Fig. 3e**). *IGHV* gene usage was similar between breast milk IgA and serum IgG repertoires (**Fig. 3f**). Breast milk IgA clonotypes had lower mean CDRH3 net charge than serum IgA clonotypes (-0.147 vs. 0.076; **Supplementary Fig. 10d**).

Despite relatively stable breast milk IgA titers, the milk antibody repertoire itself changed markedly over time. Bray-Curtis dissimilarity^34^ between Wk4 and Wk12 breast milk IgA repertoires exceeded the corresponding dissimilarity between M_D28 and M_Deliv serum IgG repertoires in all donors (**Fig. 3g**), consistent with greater longitudinal remodeling of IgA clonotypes in the milk compartment. Although this difference did not reach significance in the current cohort (*P* = 0.062), the consistent donor pattern suggests that postpartum breast milk IgA repertoires are more dynamic than the corresponding serum IgG repertoires. These data indicate that vaccine-elicited breast milk IgA is largely compartmentalized from the systemic response.

### Functional characterization of monoclonal antibodies derived from maternal serum and breast milk clonotypes

To characterize the functional properties of antibodies representative of the abundantly present maternal clonotypes, we recombinantly expressed monoclonal antibodies (mAbs) derived from maternal serum IgG and breast milk IgA clonotypes, selecting clonotypes by abundance from those with available paired V_H_:V_L_ sequences (**Fig. 4a**). Full-length heavy and light chain sequences were synthesized and expressed as human IgG1, enabling direct functional comparison across clonotypes from different compartments and vaccination timing groups. We measured S and receptor binding domain (RBD) binding and neutralization against multiple SARS-CoV-2 strains (**Fig. 4b**). Serum IgG-derived mAbs from the early group bound Wuhan S with uniformly high affinity (50% effective concentration, EC_50_ < 1 nM for all 7 mAbs), whereas late-group mAbs were more variable, with only 2 of 7 reaching that affinity threshold and several binding more weakly at 10-100 nM (**Fig. 4c**). RBD binding was detected in 4 of 7 early-group mAbs and 3 of 7 late-group mAbs; mAbs that did not bind RBD largely failed to neutralize any strain.

**Figure 4.**
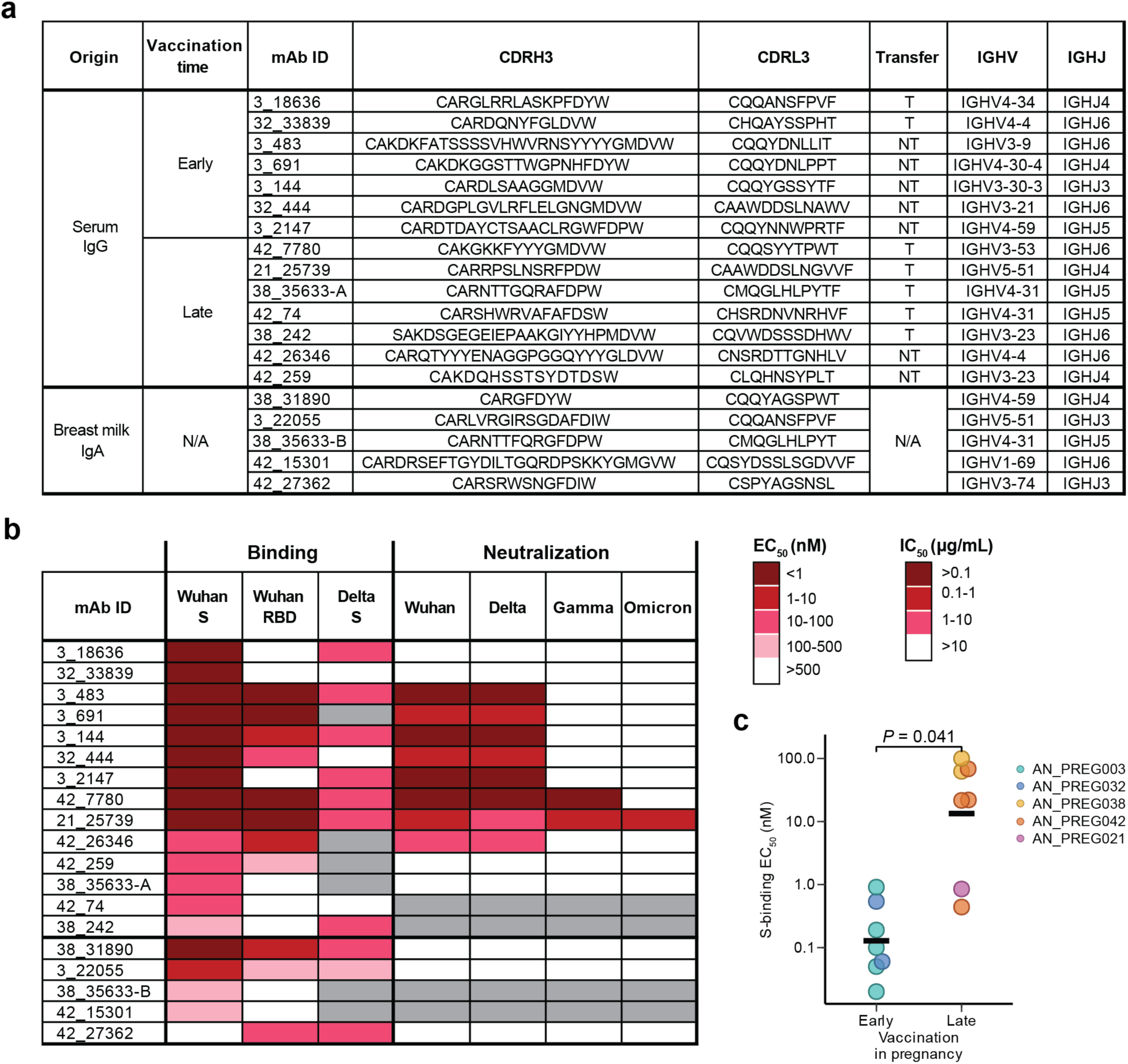
Functional properties of mAbs derived from maternal serum IgG and breast milk IgA clonotypes. **a,** Recombinantly expressed mAbs derived from early and late immunization group serum IgG clonotypes and breast milk IgA clonotypes. mAb ID indicates donor and clonotype identity. 38_35633-A and 38_35633-B belong to the same clonotype but were identified by distinct CDRH3 peptides in serum and breast milk samples, respectively, and are listed separately. Transfer classification (T, transferred; NT, non-transferred) applies to serum IgG-derived mAbs only. N/A, not applicable. **b,** Binding and neutralization profiles of expressed mAbs. Left, S- and RBD-binding (EC_50_) values measured by ELISA against Wuhan S, Wuhan RBD, and Delta S. Right, neutralization (IC_50_) values against indicated strains, tested for mAbs with Wuhan S-binding EC_50_ < 50 nM. Gray cells indicate mAbs not tested. **c,** S-binding EC_50_ values for serum IgG-derived mAbs from early and late vaccination groups. Crossbar indicates mean.

Five of seven early-group mAbs neutralized Wuhan and Delta (50% inhibitory concentration, IC_50_ < 1 µg/ml), but none neutralized Gamma or Omicron. Two late immunization group mAbs extended neutralization to three or four strains, as 42_7780 neutralized Wuhan, Delta, and Gamma (IC_50_ < 0.1 µg/mL), and 21_25739 neutralized all four strains tested. Among breast milk IgA-derived mAbs, 2 mAbs (38_31890 and 3_22055) bound Wuhan S with high affinity, comparable to the best serum IgG-derived mAbs, yet neither neutralized any strain tested. These functional data suggest that vaccination timing shapes the affinity and cross-strain neutralization of the maternal serum IgG repertoire, though the small number of expressed mAbs limits comparison between groups. Whether the high S-binding affinity of breast milk IgA-derived mAbs translates to neutralization in the native IgA format remains to be determined.

## Discussion

By integrating BCR-Seq and Ig-Seq, we resolved vaccine-elicited S-specific antibody clonotypes across matched maternal blood, cord blood, and breast milk. Our central finding is that vaccination timing imposes a trade-off with two distinct determinants shaping clonotype-level transferred immunity. Gestational age at vaccination shaped the diversity of the vaccine-elicited IgG repertoire, with earlier vaccination producing a more robust maternal serum IgG response, in terms of a greater number of clonotypes comprising the peak repertoire (**Fig. 1g**) and higher binding affinities among expressed clonotypes (**Fig. 4b,c)**. These differences may reflect the gestation-dependent modulation of humoral immunity^35^, particularly changes in B cell frequency throughout pregnancy^36^. The interval from vaccination to delivery then determined what fraction of that repertoire persisted and was available for transfer, as longer intervals between vaccination and delivery allowed greater repertoire contraction before birth, so a smaller fraction of peak clonotypes was ultimately transferred to cord. Yet those clonotypes persisting to delivery carried more somatic mutations (**Fig. 2g**) and greater intraclonal diversity (**Fig. 2i**), consistent with having undergone further affinity maturation. Conversely, later vaccination produced a more restricted IgG repertoire, with a varying range of binding affinities among the expressed mAbs, but the majority were still in circulation at delivery, yielding efficient transfer to the cord with less time for affinity maturation. Across donors, placental IgG transfer is selective at the clonotype level, in that clonotypes persisting to delivery are enriched in cord blood relative to the peak maternal repertoire.

Cord titers correlated strongest with maternal titers at M_D28 (**Fig. 2a**), whereas clonotype-level transfer was most strongly associated with maternal abundance at delivery (**Fig. 2k**). In several early-vaccinated dyads, cord titers exceeded maternal titers at delivery, a pattern consistent with prolonged transplacental accumulation over the extended period of maternal seropositivity, as has been observed across multiple vaccine antigens^17^. We interpret these observations as reflecting two distinct scales of transfer, where total fetal IgG accumulation integrates maternal IgG transfer across gestation, whereas the identity of the clonotypes transferred to cord is shaped by abundance at parturition. No single vaccination timing strategy unambiguously optimizes both the diversity and subsequent maturation (SHM and intraclonal diversity) of the peak maternal repertoire, which favors earlier vaccination, and the fraction of that repertoire and titer available at delivery for transfer, which favors later vaccination or a shorter interval to delivery. The effect of maternal vaccination timing on the quantity of transferred antibody has been examined across several vaccines, with mixed conclusions: a randomized influenza trial found cord antibody levels broadly comparable across gestational age^37^, whereas studies of SARS-CoV-2^12,23,24^ and RSV^38^ vaccination report that longer intervals between vaccination and delivery alter transfer efficiency and infant antibody levels. The present data, resolved at the clonotype level, indicate that beyond these effects on quantity, SARS-CoV-2 vaccination timing shapes which lineages are inherited and at what extent of maturation, a dimension not captured by titer alone.

Consistent with the previously reported role of IgG subclass in transplacental transfer^16,17^, our Fc analyses showed preferential transfer of IgG1 (**Supplementary Fig. 8**). Beyond subclass, the enrichment in cord blood of more somatically mutated and more intraclonally diverse clonotypes adds molecular resolution to the longstanding observation that transplacental antibody transfer is not passive or proportional^39–41^. This selectivity is consistent with the known temporal gating of placental IgG transfer, as FcRn expression on syncytiotrophoblasts increases substantially toward the third trimester^21,42^, concentrating most transplacental transfer in late gestation. Consequently, the identity of transferred clonotypes is determined primarily by which maternal IgG lineages remain abundant during that late transfer window. The finding that more mutated variants within a lineage tended to be those detected in cord is consistent with a model in which clonotypes persisting through the extended interval to delivery are those that have undergone the most affinity maturation, and that persistence to delivery is the major determinant of clonotype representation in cord. Vaccination timing therefore shapes both the clonal pool available at delivery and the properties of the clonotypes within that pool.

The breast milk compartment showed a markedly different organization. Clonal overlap between milk IgA and either serum IgG or serum IgA was minimal (**Fig. 3e**), consistent with previous proteomic analyses of human milk IgA1 repertoires^26,27^, and the few shared lineages diverged into distinct phylogenetic branches (**Fig. 3d**). This compartmentalization is notable given that vaccination was administered intramuscularly, a route that primarily engages systemic immunity, yet generated a largely distinct mucosal repertoire in milk. Shared clonotypes whose serum and milk compartment members occupied distinct, deeply diverged branches of B cell phylogenetic trees suggest that a common progenitor B cell trafficked to multiple anatomical compartments. These findings are consistent with a recent report^28^ of common and distinct origins of serum and breast milk IgA, though that study also identified a subset of shared clonotypes; the relative contribution of systemic versus mucosal priming likely depends on the route and timing of antigen exposure. Several breast milk IgA-derived mAbs had S-binding affinities comparable to the best serum IgG-derived mAbs (**Fig. 4b**), suggesting that the mucosal compartment may independently generate high-affinity S-specific antibodies through local germinal center reactions in mucosal lymphoid tissue. The breast milk IgA repertoire underwent substantial clonal remodeling between weeks 4 and 12 postpartum despite stable titers (**Fig. 3a,g**), a finding that contrasts with prior reports of high longitudinal stability in total milk IgA1 repertoires^26^. That prior work characterized total sIgA in breast milk, which is likely directed against commensal and established mucosal antigens, whereas the S-specific clonotypes tracked here represent a vaccine-elicited response that may be sustained by a more transient population seeded by systemic immunization. The remodeling we observe may therefore reflect contraction of a primary vaccine-elicited mucosal response rather than instability of the milk IgA compartment.

Several limitations of this study should be noted. The cohort of six donors, with three per vaccination timing group, limits statistical power for between-group comparisons. The small cohort size is a recognized constraint of studies requiring longitudinal sampling across multiple biological compartments, paired cord blood, and postpartum breast milk; the per-donor depth of molecular characterization was prioritized over cohort breadth. The Fc analysis was further restricted to four donors. The study was restricted to infection-naïve mothers vaccinated with BNT162b2, and all donors were enrolled from a single European clinical site, limiting generalizability across vaccine platforms, infection histories, and populations. In addition, our conclusions regarding affinity maturation are inferred from SHM, intraclonal diversity, and selected mAbs. No outcome data on infant infection or disease were available in this cohort, so the functional consequences of the clonotypic differences we describe remain to be determined. A further consideration is that maternally derived antibodies can blunt infant responses to subsequent vaccination, and the clonotype composition and maturation of transferred antibodies may influence the degree of this effect; whether the differences we observe have downstream consequences for infant immune priming is an important question for future work. Nonetheless, these data establish that passive neonatal immunity after maternal vaccination is shaped not only by how much antibody is generated but also by which clonotypes are retained, transferred, and secreted across compartments. More broadly, they suggest that optimizing maternal immunization schedules may require balancing maternal protection, placental transfer efficiency, and the molecular quality of the antibodies delivered to the infant.

## Materials and Methods

### Study subjects

All study protocols were approved by the Ethics Committee of the University of Antwerp, University Hospital of Antwerp, the Belgian Federal Agency for Medicines and Health Products (Belgium), and Dartmouth College (US). Donor PBMC and serum samples were obtained from a cohort of pregnant individuals at the Antwerp University Hospital (Belgium) under the framework of the PREGCOVAC study^43^ and secondary use within the Maternal Immunization and Determinants of Infant Immunity (MADI) Consortium, trial NCT05618548. Informed consent was obtained from the donors. A subset of 6 participants who received two doses of BNT162b2 (Comirnaty®; Pfizer-BioNTech) and had not been infected prior to dose 1 administration were analyzed in this study. Maternal blood samples collected in heparin tubes seven days after dose 2 administration were processed within 4 h of collection using SepMate^TM^ tubes to prepare PBMCs^43^. In addition, maternal blood samples collected at the time of the administration of dose 2, twenty-eight days after dose 2, and delivery, and umbilical cord blood samples at the time of delivery were processed to obtain serum. Breast milk samples were collected postpartum at week 4 and week 12 and processed within 24 h for storage at -80°C^43^. Participant metadata is available in **Supplementary Table 1**. Breast milk collection metadata is available in **Supplementary Table 5**.

### Library preparation for high-throughput sequencing of V_H_

Total RNA (500 ng) extracted from PBMCs taken 7 days after vaccination with dose 2 was used for library preparation. Reverse transcription was performed using SuperScript IV (Invitrogen) and Oligo(dT) primer (Invitrogen) according to manufacturer instructions. V_H_-specific transcripts were amplified using the FastStart High Fidelity PCR System (Roche) with gene-specific primers^30^. Single-chain amplicons were barcoded using the Next^®^ Ultra^TM^ II DNA Library Prep Kit (NEB, E7645) and sequenced on the Illumina NextSeq platform.

### V_H_:V_L_-paired sequencing

Paired heavy- and light-chain sequencing of single B cells from day 7 post-booster samples were performed, as previously described^30,44^. Briefly, single B cells were encapsulated in emulsion droplets using a custom flow-focusing microfluidic device. Droplets contained lysis buffer and poly(dT)-conjugated magnetic beads to capture mRNA transcripts encoding immunoglobulin heavy and light chains. Magnetic beads were subsequently recovered and re-emulsified to serve as templates for emulsion overlap-extension RT–PCR. Nested PCR amplification generated ∼850-bp linked V_H_:V_L_ amplicons, which were sequenced on a PacBio Sequel IIe platform.

### Purification of total IgG from serum and subsequent mass spectrometry sample preparation

For each serum sample, IgG was purified using Protein G affinity columns. Briefly, about 2 mL of the diluted serum (1:2 dilution in PBS) was filtered using 0.22-μm syringe filters and passed through 2-mL Protein G Plus agarose (Thermo Scientific) affinity column using gravity flow. The flowthrough was collected and passed through the column three times. The column was washed with 10 column volumes (CV) of PBS and eluted with 5 CV of 100 mM glycine-HCl, pH 2.7. The eluate was immediately neutralized with 2 mL of 1M Tris-HCl, pH 8.0.

SARS-CoV-2 Spike (HexaPro)^45^ was purified and immobilized on N-hydroxysuccinimide (NHS)-activated agarose resins (Pierce) by overnight rotation at 4°C. Each purified IgG sample was rotated with immobilized Spike for 2 h at RT. After incubation, the mixture was applied to a Pierce spin column (Thermo Scientific) and washed with 0.5 mL of PBS three times, resuspended in 0.2 mL of PBS and 5 μg of recombinantly expressed IdeS was added for the on-column digest of IgG into F(ab’)_2_, followed by rotation for 4 h at 37°C. After collecting flowthrough (S-specific Fc), the mixture was washed with 10 CV of PBS three times and S-enriched F(ab’)_2_ was eluted using 1% formic acid in 0.5-mL fractions and neutralized with NaOH-Tris. Elution fractions were pooled and concentrated under vacuum to a volume of ∼0.1 mL.

For each enrichment, elution and flowthrough samples were prepared for mass spectrometry. In addition, S-specific Fc samples were prepared for mass spectrometry. Briefly, samples were denatured in 50% (vol/vol) 2,2,2-trifluoroethanol (TFE), 50 mM ammonium bicarbonate and 10 mM dithiothreitol (DTT) at 55°C for 45 min, then alkylated by incubation with 32 mM iodoacetamide (Sigma) for 30 min at RT in the dark. Alkylation was quenched by the addition of 20 mM DTT. Samples were diluted tenfold with 50 mM ammonium bicarbonate and digested with either trypsin (Promega) for F(ab’)_2_ samples or Lys-C (Promega) for Fc samples (1:30 enzyme:protein) for 16 h at 37°C. Formic acid was added to 1% (vol/vol) to quench the digestion, and the sample volume was reduced to ∼100 μL under vacuum. Peptides were then purified using C18 spin columns (Thermo Scientific, 89870), washed three times with 0.1% formic acid, and eluted with a 60% acetonitrile and 0.1% formic acid solution. C18 eluate was concentrated under vacuum centrifugation and resuspended in 50 μL in 5% acetonitrile, 0.1% formic acid.

### Purification of total IgA from serum and breast milk and subsequent mass spectrometry sample preparation

Breast milk was processed for delipidation and removal of cells by centrifugation at 3000xg at 4°C, isolation of the middle skim layer and second centrifugation 3000xg at 4°C. The skim layer was used for IgA purification using Peptide M resin column. Each serum sample flowthrough from the initial Protein G and breast milk skim layer was used for IgA purification using Peptide M resin column according to manufacturer instructions. Purified IgA was processed similarly to serum IgG for enrichment for Spike. However, no cleavage to F(ab’)_2_ and Fc fragments was performed and full-length IgA samples were prepared for mass-spectrometry.

### LC–MS/MS analysis

Samples were injected into the Thermo Orbitrap Fusion equipped with an electron transfer dissociation(ETD) module and analyzed using separate acquisition methods for F(ab’)_2_ and Fc.

For the F(ab’)_2_ analysis, the following LC gradient was used, all at 300 nL/min: 1.6% acetonitrile for 5 minutes, 1.6-9.6% acetonitrile over 5 minutes, 9.6-25.6% acetonitrile over 50 minutes, 25.6-32% acetonitrile over 10 minutes, 32%-76% acetonitrile over 2 minutes, and 76% acetonitrile for 10 minutes. The mass spectrometer was run in positive mode, with the MS1 scan detected in the orbitrap at a resolution of 120,000. The scan range was limited to 375-1600 m/z and the radio frequency (RF) lens was set to 60%. The normalized automatic gain control (AGC) target was set to 125%.

Monoisotopic peak determination was used on the peptide level and an intensity threshold of 1E4 was used. Only determined charge states between 2-6 were used and dynamic exclusion was applied after 2 detections (in a 10 ppm tolerance) within 30 seconds for a duration of 15 seconds. MS2 scans were detected in the ion trap and fragmented with 30% higher-energy collisional dissociation (HCD) energy in rapid mode. The normalized AGC target was set to 200%.

For Fc analysis, the following LC gradient was used, all at 300 nL/min: 4.8-20.8% acetonitrile over 40 minutes, 20.8-25.6% acetonitrile over 40 minutes, 25.6-40% acetonitrile over 30 minutes, 40%-80% acetonitrile over 6 minutes, 80% acetonitrile for 4 minutes, 80-4.8% acetonitrile over 1 minute, and 4.8% acetonitrile for 9 minutes. The mass spectrometer was run in positive mode, with the MS1 scan detected in the orbitrap at a resolution of 120,000. The scan range was limited to 350-1800 m/z and the RF lens was set to 60%. Monoisotopic peak determination was used on the peptide level and only determined charge states between 2-8 are used. Dynamic exclusion was applied after 1 detection (in a 10 ppm tolerance) for a duration of 20 seconds. An intensity filter of 5E4 was used. An MS2 scan in the orbitrap, at a resolution of 30000, was run with an isolation window of 2 m/z using HCD collision at 28%. If a m/z of 204.0867 (HexNac), 138.0545 (HexNac fragment), 366.1396 (HexNacHex), 163.081 (Hex), 292.103 (NeuAc), or 308.098 (NeuGc) were detected within a 15 ppm tolerance further scans were initiated: A MS2 orbitrap HCD scan, with a stepped collision energy of 20, 30, and 40% at a resolution of 30000 and an orbitrap electron transfer combined with higher-energy collision dissociation (EThcD), with a supplemental 15% collision energy and resolution of 30,000.

### MS/MS data analysis

Donor-specific peptide search databases for MS data acquisition were created using V_H_ sequences from high-throughput sequencing of V_H_ and V_L_ repertoires of single B cells from dose 2 day 7 donor samples, as previously described^31,32^. V_H_ amino acid sequences with ≥2 reads from high-throughput sequencing were included to construct the peptide search database. These V_H_ sequences were then combined with a database of background proteins, which included a consensus human protein database (Ensembl 73, longest sequence/gene), and a list of common protein contaminants (MaxQuant). Spectra were searched against the database using SEQUEST (Proteome Discoverer 2.4; Thermo Scientific). For IgA searches, the database was modified by substituting the IgG CH1 N-terminal tryptic sequence ASTK with the IgA CH1 equivalent ASPTSPK to enable correct assignment of IgA-derived peptides. Searches considered fully tryptic peptides only, allowing up to two missed cleavages. A precursor mass tolerance of 5 ppm and fragment mass tolerance of 0.5 Da were used. Modifications of carbamidomethyl cysteine (static), oxidized methionine, and formylated lysine, serine or threonine (dynamic) were selected. High-confidence peptide-spectrum matches (PSMs) were filtered at a false discovery rate of <1% as calculated by Percolator (q-value < 0.01, Proteome Discoverer 2.4; Thermo Scientific). Iso/Leu sequence variants were collapsed into single peptide groups. For each scan, PSMs were ranked first by posterior error probability (PEP), then q-value, and finally cross-correlation (XCorr) score. Only unambiguous top-ranked PSMs were kept; scans with multiple top-ranked PSMs (equivalent PEP, q-value, and XCorr) were designated ambiguous identifications and removed. The average mass deviation (AMD) for each peptide was calculated as previously described^46^. Peptides with AMD > 1.5 ppm were removed. Peptide abundance was calculated from the extracted-ion chromatogram (XIC) peak area, as previously described^46^. For each peptide, a total XIC area was calculated as the sum of all unique peptide XIC areas of associated precursor ions. The average XIC area across replicate injections was calculated for each sample. For each dataset, the eluate and flowthrough abundances were compared, and serum abundance of Spike-specific antibody species was estimated by retaining peptides with high confidence of antigen specificity, indicated by an eluate with at least a 5-fold higher XIC signal.

### Clonotype indexing and peptide-to-clonotype mapping

High-confidence peptides identified through MS/MS analysis were mapped to clonotype clusters. Peptides that are uniquely mapped to a single clonotype were considered ‘informative’, and clonotypes detected ≥ 2 PSM were kept for as high-confidence identifications. The abundance of each antibody clonotype was calculated by summing the XIC areas of the informative peptides mapping to ≥ 4 amino acids of the CDRH3 region. Shared antibody clonotypes across serum IgG, IgA and breast milk IgA were defined as the ones mapping to the same ClusterID from the donor-specific search databases.

### Quantifying relative abundances of antibody clonotypes

High-confidence peptides identified by MS/MS were mapped to antibody clonotypes. Peptides that uniquely mapped to a single clonotype were considered informative. The abundance of each clonotype was calculated by summing the XIC areas of informative peptides that mapped to ≥4 amino acids within the CDRH3 region. The relative amount of serum IgG was estimated by multiplying the clonotype’s fraction of the serological repertoire (%) by the binding titer (1/ED_50_) of the corresponding serum sample against Spike, as previously described^31,33,47^.

### Lineage analysis

B cell lineage analysis was conducted using the Immcantation software suite. V_H_-only BCR sequencing reads were merged using PEAR^48^ v0.9.6. Next, low quality reads were removed, primer regions were masked, duplicate reads were collapsed, and only sequences with at least 2 reads were retained using scripts from the pRESTO^49^ package v0.7.9. Sequences were annotated using IgBLAST^50^ v1.22.0 using the IMGT germline database^51^ accessed on May 5, 2024. Then Adaptive Immune Receptor Repertoire (AIRR)-formatted databases were generated, and productive heavy chain sequences were retained using Change-O^52^ package v1.3.4.

Paired-chain PacBio sequencing reads were filtered for high quality reads, then primer identification was used to determine the heavy and light chain portions of each read using pRESTO. Duplicate sequences were collapsed, then the heavy and light chain portions were annotated using IgBLAST, and AIRR-formatted databases were made for heavy and light chain sequences respectively using Change-O. Productive sequences matching the respective locus (IGH vs IGK/L) were retained, then the heavy and light chain databases were merged.

The V_H_-only and paired-chain sequence databases were merged, then novel alleles were detected and genotypes inferred using TIgGER^52,53^ v1.1.0. To remove sequences with partial V genes, sequences with more than 3 ambiguous V-gene calls were removed. Using Shazam^52^ v1.2.0, we automatically detected the clonal threshold using the length-normalized Hamming distance. Clonal clustering was performed using single-linkage hierarchical clustering using Scoper^54^ package v1.3.0. Phylogenetic trees were constructed using Dowser^55,56^ v2.4.1.999 implementing RaxML^57^ (using “scaled” partition on donors with V_H_:V_L_ data) using R v4.5.3 and visualized with ggtree^58^ v3.16.0. Large clones were downsampled with ‘sampleClones’ weighted by duplicate count with even sampling across Ig-seq match groups prior to visualization.

### Somatic hypermutation level analysis

SHM levels for serum clonotypes were estimated from B cell sequences encoding the corresponding serum CDRH3 peptides identified Ig-Seq. Mutation frequencies calculated by the ‘observedMutations’ function in Shazam within IMGT_V region were extracted from class-switched (IgG or IgA) B cell sequences matching the CDRH3 peptides. Mutation rates were first averaged within each B cell clone, then averaged across clones corresponding to the serum clonotype. When only low confidence sequence matches were available, the clonotype was excluded from downstream analysis and statistical modeling.

### Generalized linear mixed-effects model (GLMM) of Fab features

We implemented the ‘glmer’ function of lme4^59^ v1.1-34 package in R v4.5.0 with formula *Transfer ∼ M_D0 + M_D28 + M_Deliv + SHM + intraclonal_diversity + CDRH3_hydrophobicity + CDRH3_netcharge + CDRH3_length + (1|donor)*. Clonotypes with missing SHM values were excluded from this analysis.

The amount of IgG at each time point was determined by multiplying the relative abundance of the clonotype by the corresponding S-reactive titer for that sample. The output variable, *Transfer*, was defined as a binary indicator for clonotype detection in cord blood. *CDRH3_hydrophobicity*, *CDRH3_netcharge*, and the *CDRH3_length* of each clonotype were calculated using the Peptides^60^ v2.4.6 package ‘hydrophobicity’ function with Kyte-Doolittle scale^61^, ‘charg’ function at pH 7 with Lehninger pK scale, and ‘length’ function, respectively. The model plot was generated using sjPlot^62^ v2.8.26 and ggplot2^63^ v3.5.1. Model robustness was assessed by leave-one-out analysis, excluding one donor each time. Multicollinearity among predictors was assessed using variance inflation factor (VIF) with car^64^ package v3.1.3.

### Fc subclass-glycoform identification and quantification and statistical analysis

Fc samples were searched using a database of expected Fc sequences from IMGT GENE-DB^65^ using pGlyco3 software^66^. Formylation was allowed as a dynamic modification of lysine, serine, and threonine, oxidation was included as a dynamic modification on methionine, dynamic acetyl and methionine losses were allowed on the N-terminus, and carbamidomethyl modifications were static on the C-terminus. Up to 2 missed cleavages were allowed and the precursor tolerance was set to 10 ppm and the fragment tolerance was set to 20 ppm. The abundance of each subclass-glycoform was calculated by summing the isotope area of total detected glycoforms within a subclass with Gly Score >3.

To perform statistical testing on the clustering of the Fc glycan profiles, we used the pairwise Euclidean distance matrix, calculated on glycan/sample profiles, as input to the PERMANOVA^67^ test using the R package vegan^68^ v2.7.1 function ‘adonis2’ with formula *Distance_matrix ∼ timepoint + donor*, reporting the marginal effects of each term. The analysis was performed separately on the samples and the glycans. For the glycans, the formula *Distance_glycans ∼ subclass + bisection + fucosylation + sialylation + galactosylation* was used.

For the Fc linear mixed-effects model, we used the ‘lmer’ function of lme4^59^ v1.1-34 with formula *Cord ∼ 0 + M_D0 + M_D28 + M_Deliv + subclass + fucosylation + sialylation + galactosylation + bisection + (1|donor)*. We modeled using an LMM instead of a GLMM because many Fc glycans were also detected in cord, so the outcome predictor was amount of transfer. The model was restricted to S-specific samples. The outcome variable, *Cord*, was the amount of each glycan within each IgG subclass in cord, calculated using the relative abundance multiplied by the S-reactive titer for the respective sample. Similar calculations were performed for the amounts in maternal serum. *Fucosylation*, *sialylation*, *galactosylation*, and *bisection* were implemented as binary indicators of sugar presence in the glycan. Plots were generated using sjPlot^62^ v2.8.26 and ggplot2^63^ v3.5.1. Model robustness was assessed by leave-one-out analysis, excluding one donor each time.

### Recombinant antibody expression and purification

Selection of antibody sequences corresponding to serum IgG and IgA clonotypes for recombinant expression was guided by clonally assigned B cell sequences together with proteomic data. We selected V_H_:V_L_-paired sequences corresponding to detected serum CDRH3 peptides. When no exact V_H_:V_L_-paired match was available, the sequence with the most similar CDRH3 was selected. Among candidates, sequences with the highest expansion (highest duplicate read count) were prioritized. Full-length heavy and light chain sequences were then synthesized in human IgG1 expression vectors by Twist Bioscience. Each mAb was either produced by Twist’s IgG production service or transfected in-house into Expi293 cells with PEI at a 1:3 heavy to light chain DNA ratio. After 6 days of incubation at 37°C with 8% CO_2_, the supernatants were collected. IgG was purified by passing through protein G agarose resin (Thermo Scientific, 20397). The column was washed with 20 mL of PBS, antibodies were eluted with 5 mL 100 mM glycine-HCl, pH 2.7, and the eluate was immediately neutralized with 0.75 mL 1 M Tris-HCl, pH 8.0. Antibodies were then buffer exchanged into PBS using Amicon Ultra-30K centrifugal filter (Millipore).

### ELISA

ELISA plates (Corning 3361, Thermo Fisher Scientific) were coated with SARS-CoV-2 S Hexapro protein or Wuhan Hu-1 RBD protein^45^ at 4 μg/mL in PBS for 50 μL per well and incubated overnight at 4°C. Plates were washed three times with washing buffer (PBST; 1X PBS containing 0.05% Tween-20 (RPI)) and blocked with 200 μL blocking buffer (PBSM; 1X PBS containing 0.5% BSA (VWR)) for 2 h at RT. Monoclonal antibodies were diluted in PBSM to the desired starting concentration and applied to plates by serial dilution, followed by 1 h incubation at RT. Plates were then washed three times with PBST, and HRP-conjugated anti-human IgG secondary antibody (A0293, Sigma Aldrich) diluted 1:5,000 in PBSM was added and incubated for 40 mins at room temperature. Following incubation, plates were washed three times with PBST. Bound antibodies were detected using 50 μL of 3,3′,5,5′-tetramethylbenzidine (TMB) substrate (Thermo Fisher Scientific) and developed for 5 mins. The reaction was stopped by adding 100 μL of 2 N H_2_SO_4_ (VWR). Absorbance was measured at 450 nm using SpectraMax Paradigm microplate reader (Molecular Devices) with SoftMax Pro v7.0.2. Data were analyzed and fitted for EC_50_ using a 4-parameter logistic nonlinear regression model in the GraphPad Prism 10.6.1 software.

### Cell lines

HEK 293T/17 (ATCC CRL-11268) and HEK 293T/ACE2 (provided by Dr. Michael Farzan, The Scripps Institute) cells were cultured in DMEM (Gibco) containing 10% fetal bovine serum (GeminiBio) and penicillin/streptomycin (Gibco). Puromycin (3 μg/mL, Gibco) was added to HEK 293T/ACE2 cell growth medium to maintain the expression of ACE2 receptor.

### Pseudovirus production

Pseudoviruses were produced by co-transfection of HEK-293T/17 cells with plasmids encoding a lentivirus backbone vector (pCMV ΔR8.2)^69^, firefly luciferase reporter gene (pHR’ CMV Luc)^69^, a human transmembrane protease serine 2 (TMPRSS2)^70^, and a full-length SARS-CoV-2 spike from an early isolate (B.1.1), Delta (B.1.617.2), Gamma (P.1), or Omicron (BA.2.75.2) using FuGENE 6 transfection reagent (Promega). Between 48 and 72 hours following transfection, supernatants containing pseudoviruses were harvested, filtered with a 0.45-μm Steriflip vacuum tube (EMD Millipore), and stored at -80°C until use. Tissue culture infectious dose 50 (TCID_50_) was determined by the dilution producing ∼200,000 relative luminescence units (RLU) in the absence of serum that was used for the neutralization assay.

### Neutralization assays

Neutralization titers of serum IgG, breast milk, and isolated mAb were obtained using a lentivirus-based assay and calculated as functions of the reduction in luciferase gene expression following a single round of infection with SARS-CoV-2 spike-pseudotyped viruses in ACE2 overexpressing HEK293T/ACE cell line, as described^71^. Pseuduoviruses were co-incubated with eight serial five-fold dilutions of heat inactivated breast milk, enriched IgG from serum, or monoclonal antibodies (10 – 0.0032 µg/mL) in duplicate in PLL cell culture-treated 96-well plates for 45 min at 37°C, prior to the addition of HEK293T/ACE2 cells (10^4^ cells/well). Following incubation for 72 h at 37°C and 5% CO_2_, cells were disrupted on a platform shaker at 600 rpm in Luciferase cell culture lysis reagent (Promega). Luciferase was activated using the Bright-Glo Luciferase Assay System and the RLUs were measured on a Spectramax L Luminometer (Molecular Devices). Serum IgG and breast milk neutralization titers were calculated as the inhibitory dilution of serum at which RLU are determined to be 50% (ID_50_) compared to virus-only control wells after subtracting cell-only background luminescence. mAb neutralization titers were calculated as the inhibitory concentration of antibody at which RLU are determined to be 50% (IC_50_) compared to virus-only control wells after subtracting cell-only background luminescence.

### Use of large language models

Large language models, including Claude and ChatGPT, were used in the development of this manuscript to assist with debugging code and editing of the manuscript. All AI-generated outputs were reviewed and adapted before using in the context of this study.

### Statistics

All statistical comparisons are calculated using two-tailed Wilcoxon rank-sum (Wilcoxon signed-rank when paired) test at ɑ = 0.05 unless reported otherwise. When relevant, multiple hypothesis correction was performed using the Benjamini-Hochberg correction. All correlations reported are Spearman correlations unless otherwise specified. Statistics were calculated using ggpubr^72^ v0.6.0 and rstatix^73^ v0.7.2. Plots were generated using ggplot2^63^ v3.5.1, ggbeeswarm^74^ v0.7.3, ggh4x^75^ v0.3.1, ggalluvial^76^ v0.12.5, and ComplexHeatmap^77^ v2.24.0.

## Data availability

The proteomics data reported in this paper are archived through MassIVE (https://massive.ucsd.edu/ProteoSAFe/static/massive.jsp) under accession code MSV000101060.

## Acknowledgements

This work was supported in part by the National Institute of Allergy and Infectious Diseases (U19AI145825), the National Institute of General Medical Science (P20GM113132) and by a Korea University Grant. H.J.M. was supported by National Cancer Institute grant T32CA134286.

## Author contributions

A.M., M.E.A., and J.L. conceived the project and supervised laboratory analyses. A.Y. and L.P. led the data generation and analysis. N.C.C, N.M.T, and M.R.S. assisted with proteomic experimentation and analysis. T.K. provided neutralization data. J.Y., E.O., E.V., T.Y., B.K., D.N.K., and M.C. assisted with experiments and analysis. H.J.M. and K.B.H. supervised bioinformatic and statistical analysis. L.D.B. and K.M. contributed to donor enrollment and clinical sample collection and management. P.P. helped with the management and logistics of the clinical samples. A.Y., L.P., and J.L. wrote the manuscript with input from co-authors. All authors reviewed and edited the manuscript.

## Declaration of interests

K.B.H. has received consulting fees from Prellis Biologics. All other authors declare no competing interests.

---

**Supplementary Table 1.** Cohort donor metadata.

| Donor ID | Age during pregnancy | Ethnicity | Healthcare worker? | Gestational age (Wk) |  |  | Days between Dose 2 to Delivery | Infant sex |
| --- | --- | --- | --- | --- | --- | --- | --- | --- |
|  |  |  |  | Dose 1 | Dose 2 | Delivery |  |  |
| AN_PREG003 | 25 | Caucasian | Yes | 14.1 | 18.0 | 39.1 | 148 | Female |
| AN_PREG032 | 35 | Caucasian | Yes | 14.7 | 19.7 | 41.3 | 151 | Male |
| AN_PREG001 | 34 | Caucasian | Yes | 17.3 | 20.4 | 40.1 | 138 | Male |
| AN_PREG038 | 29 | Caucasian | No | 26.4 | 31.4 | 40.4 | 63 | Female |
| AN_PREG042 | 29 | Caucasian | No | 28.1 | 33.1 | 40.6 | 53 | Female |
| AN_PREG021 | 30 | Caucasian | Yes | 32.3 | 35.3 | 40.6 | 37 | Female |

**Supplementary Table 2.** BCR-Seq summary.

|  |  |  | AN_PREG003 | AN_PREG032 | AN_PREG001 | AN_PREG038 | AN_PREG042 | AN_PREG021 |
| --- | --- | --- | --- | --- | --- | --- | --- | --- |
| Single chain (Illumina NextSeq) | V<sub>H</sub> class-switched (IgG + IgA) | Raw paired reads | 2,820,690 | 3,499,230 | 4,535,396 | 2,328,527 | 2,980,309 | 2,696,073 |
|  |  | Unique sequences | 687,066 | 1,276,148 | 1,769,208 | 741,462 | 1,623,760 | 960,786 |
|  | V<sub>H</sub> non-class-switched (IgM) | Raw paired reads | 2,408,473 | 1,880,251 | 2,829,197 | 2,396,866 | 3,540,987 | 2,962,349 |
|  |  | Unique sequences | 650,121 | 696,733 | 1,216,037 | 847,876 | 1,712,647 | 795,601 |
| Paired chain (PacBio Sequel IIe) | V<sub>H</sub>:V<sub>L</sub> class-switched (IgG + IgA) | Raw consensus reads | 572,638 | 422,990 | N/A | 396,865 | 733,397 | 518,764 |
|  |  | Unique sequences | 514,082 | 389,340 | N/A | 350,242 | 652,567 | 494,298 |
|  | V<sub>H</sub>:V<sub>L</sub> non-class-switched (IgM) | Raw consensus reads | 252,274 | 418,638 | N/A | 327,571 | 688,757 | 590,177 |
|  |  | Unique sequences | 216,185 | 377,053 | N/A | 273,237 | 598,272 | 564,628 |

**Supplementary Table 3.**
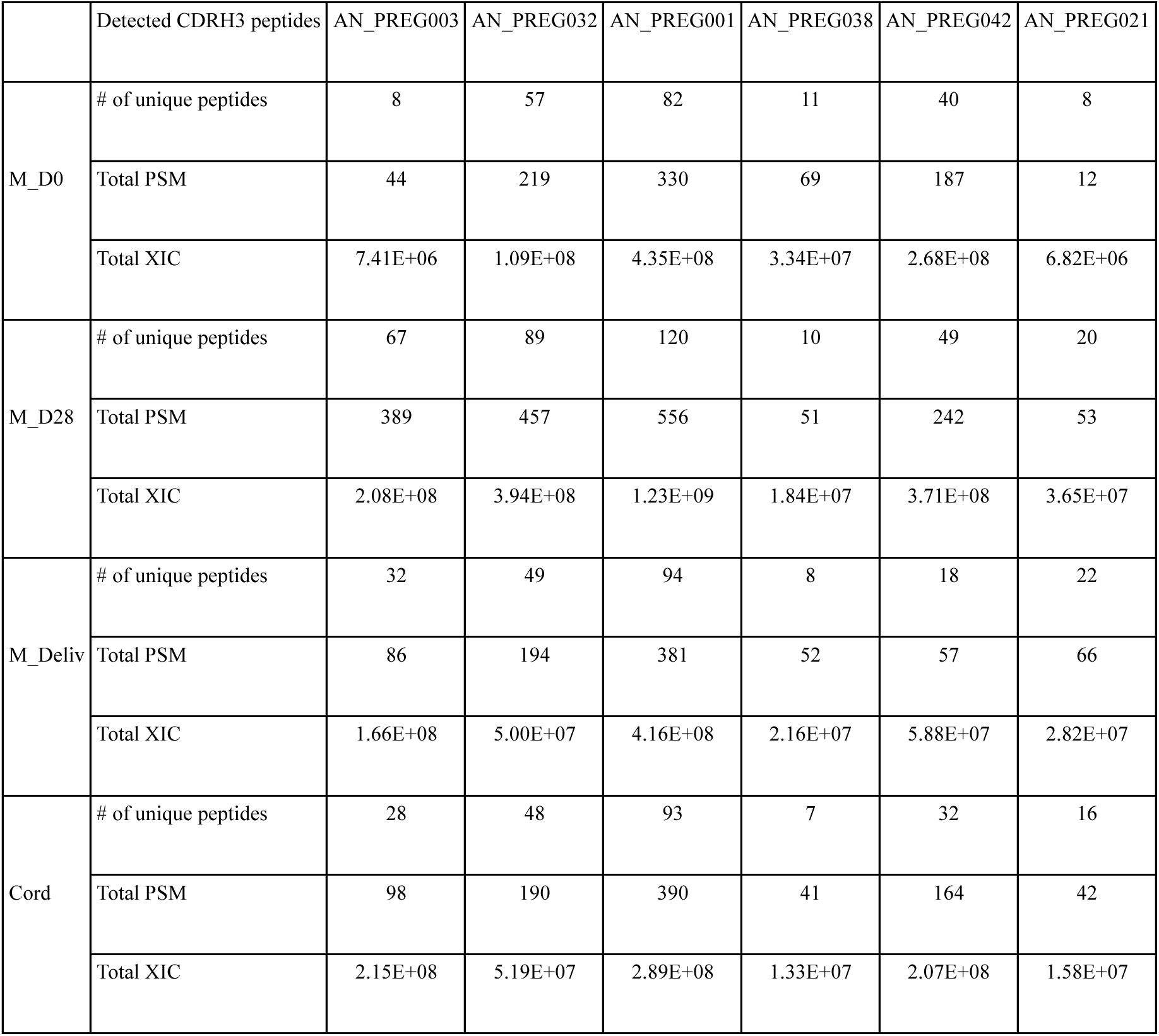
Ig-Seq summary.

|  | Detected CDRH3 peptides | AN_PREG003 | AN_PREG032 | AN_PREG001 | AN_PREG038 | AN_PREG042 | AN_PREG021 |
| --- | --- | --- | --- | --- | --- | --- | --- |
| M_D0 | # of unique peptides | 8 | 57 | 82 | 11 | 40 | 8 |
|  | Total PSM | 44 | 219 | 330 | 69 | 187 | 12 |
|  | Total XIC | 7.41E+06 | 1.09E+08 | 4.35E+08 | 3.34E+07 | 2.68E+08 | 6.82E+06 |
| M_D28 | # of unique peptides | 67 | 89 | 120 | 10 | 49 | 20 |
|  | Total PSM | 389 | 457 | 556 | 51 | 242 | 53 |
|  | Total XIC | 2.08E+08 | 3.94E+08 | 1.23E+09 | 1.84E+07 | 3.71E+08 | 3.65E+07 |
| M_Deliv | # of unique peptides | 32 | 49 | 94 | 8 | 18 | 22 |
|  | Total PSM | 86 | 194 | 381 | 52 | 57 | 66 |
|  | Total XIC | 1.66E+08 | 5.00E+07 | 4.16E+08 | 2.16E+07 | 5.88E+07 | 2.82E+07 |
| Cord | # of unique peptides | 28 | 48 | 93 | 7 | 32 | 16 |
|  | Total PSM | 98 | 190 | 390 | 41 | 164 | 42 |
|  | Total XIC | 2.15E+08 | 5.19E+07 | 2.89E+08 | 1.33E+07 | 2.07E+08 | 1.58E+07 |

**Supplementary Table 4.**
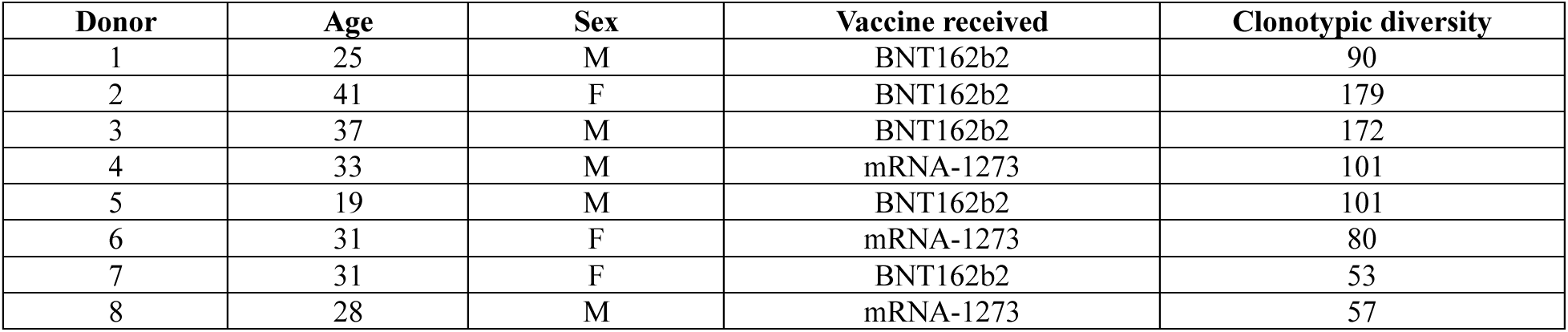
Non-pregnant donor metadata.

| Donor | Age | Sex | Vaccine received | Clonotypic diversity |
| --- | --- | --- | --- | --- |
| 1 | 25 | M | BNT162b2 | 90 |
| 2 | 41 | F | BNT162b2 | 179 |
| 3 | 37 | M | BNT162b2 | 172 |
| 4 | 33 | M | mRNA-1273 | 101 |
| 5 | 19 | M | BNT162b2 | 101 |
| 6 | 31 | F | mRNA-1273 | 80 |
| 7 | 31 | F | BNT162b2 | 53 |
| 8 | 28 | M | mRNA-1273 | 57 |

**Supplementary Table 5.**
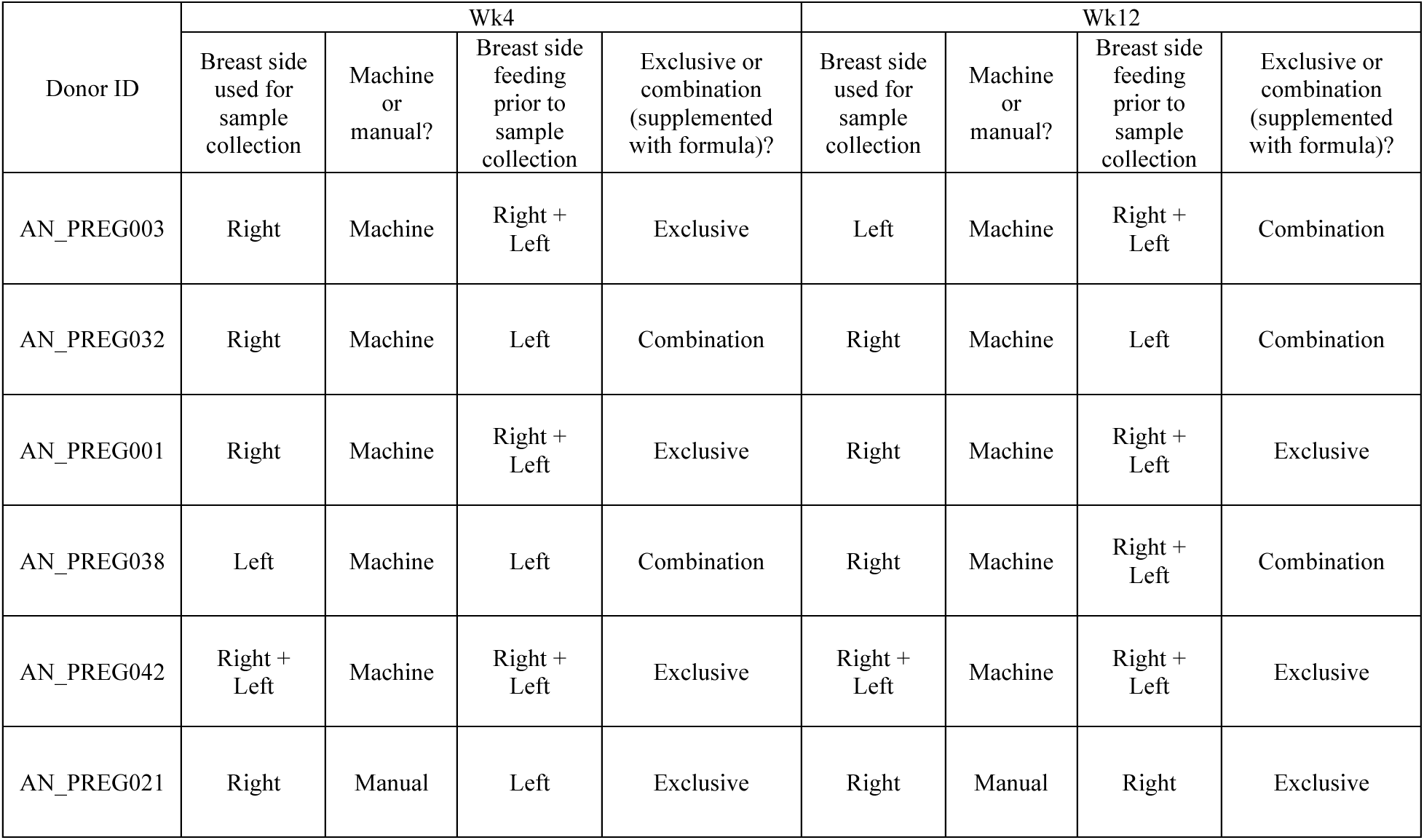
Breast milk sample collection metadata.

| Donor ID | Wk4 |  |  |  | Wk12 |  |  |  |
| --- | --- | --- | --- | --- | --- | --- | --- | --- |
|  | Breast side used for sample collection | Machine or manual? | Breast side feeding prior to sample collection | Exclusive or combination (supplemented with formula)? | Breast side used for sample collection | Machine or manual? | Breast side feeding prior to sample collection | Exclusive or combination (supplemented with formula)? |
| AN_PREG003 | Right | Machine | Right + Left | Exclusive | Left | Machine | Right + Left | Combination |
| AN_PREG032 | Right | Machine | Left | Combination | Right | Machine | Left | Combination |
| AN_PREG001 | Right | Machine | Right + Left | Exclusive | Right | Machine | Right + Left | Exclusive |
| AN_PREG038 | Left | Machine | Left | Combination | Right | Machine | Right + Left | Combination |
| AN_PREG042 | Right + Left | Machine | Right + Left | Exclusive | Right + Left | Machine | Right + Left | Exclusive |
| AN_PREG021 | Right | Manual | Left | Exclusive | Right | Manual | Right | Exclusive |

**Supplementary Figure 1.**
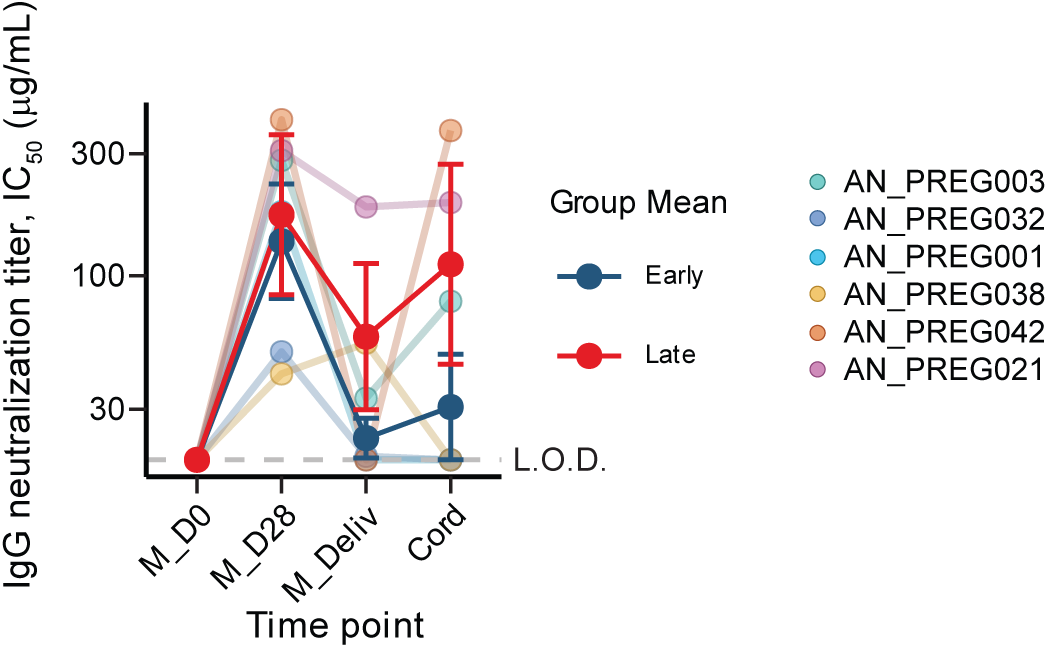
Additional characterization of maternal and cord serum samples. Maternal and cord S-reactive IgG neutralizing titers at the indicated time points in pregnancy and in cord. Thin lines represent individual donors, and thick lines represent group means, with error bars indicating s.e.m.

**Supplementary Figure 2.**
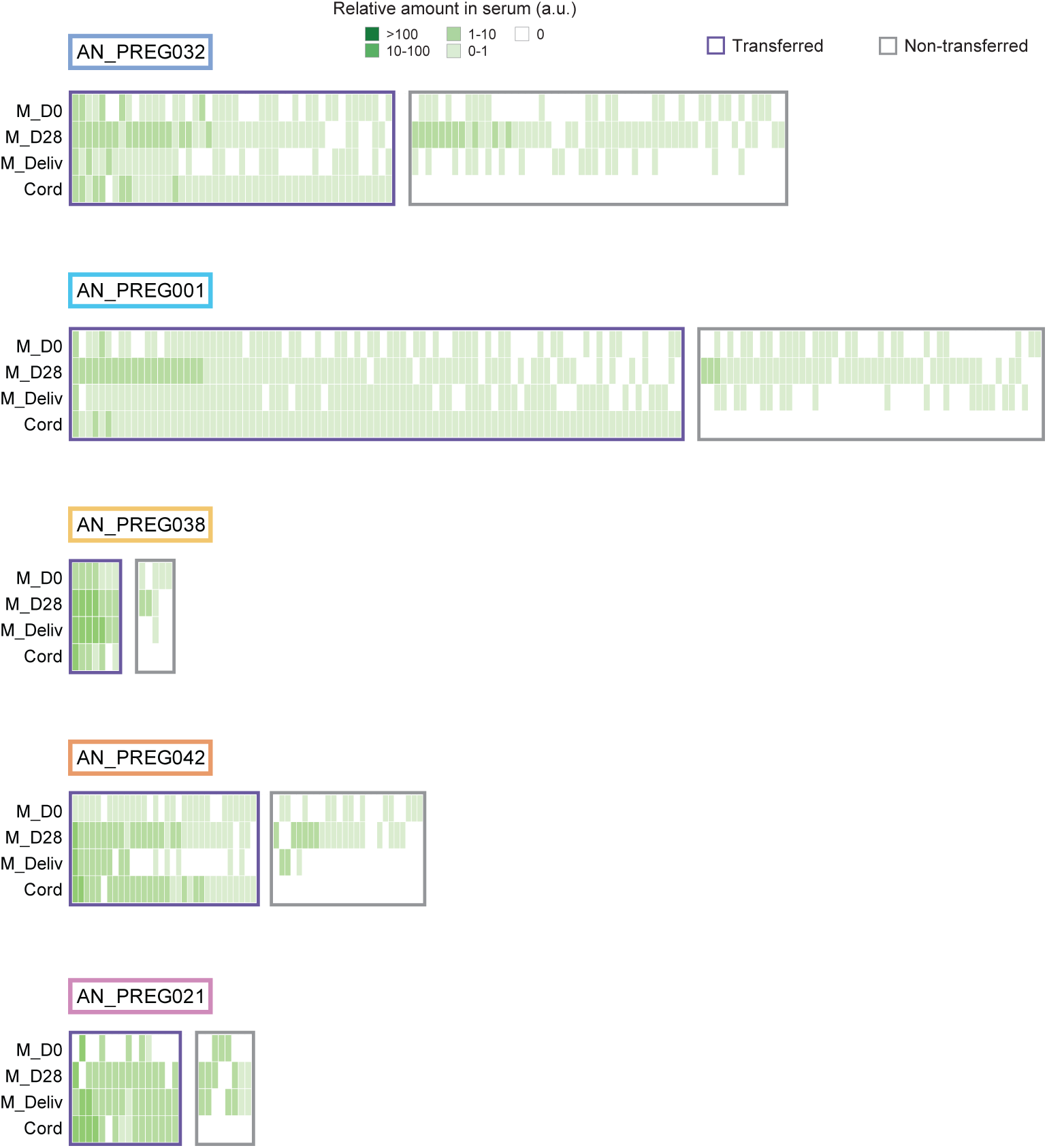
Relative serum IgG clonotype abundance heatmaps from five donors. Each column represents an individual IgG clonotype, with color indicating its relative amount in serum (arbitrary units), calculated by scaling the anti-S serum IgG binding titer by the relative abundance of each clonotype determined by proteomic analysis. Clonotypes were classified as transferred (detected in cord) or non-transferred (absent in cord).

**Supplementary Figure 3.**
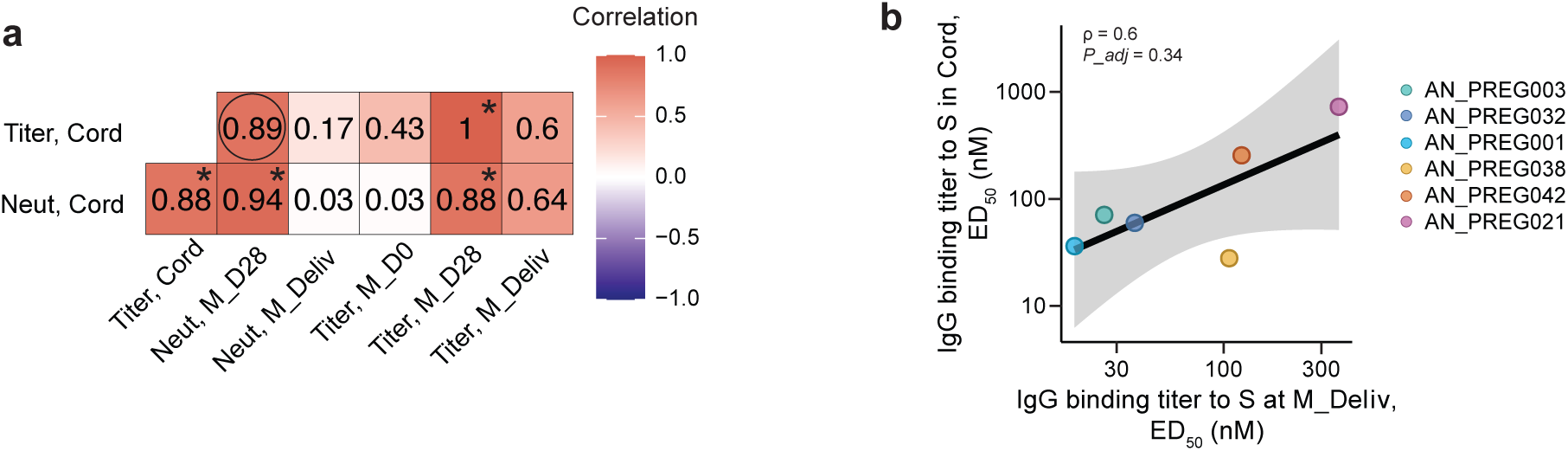
Additional characterization of maternal serological IgG in the context of transfer to cord. **a,** Correlation matrix for S-reactive IgG binding and neutralization titers across maternal serum time points and cord blood. The asterisk indicates a statistically significant correlation (*P* < 0.05 after Benjamini-Hochberg correction) and the circle represents *P* < 0.1. **b,** Correlation between S-specific IgG binding titer in paired maternal serum and cord at delivery. The regression line is shown with 95% confidence interval.

**Supplementary Figure 4.**
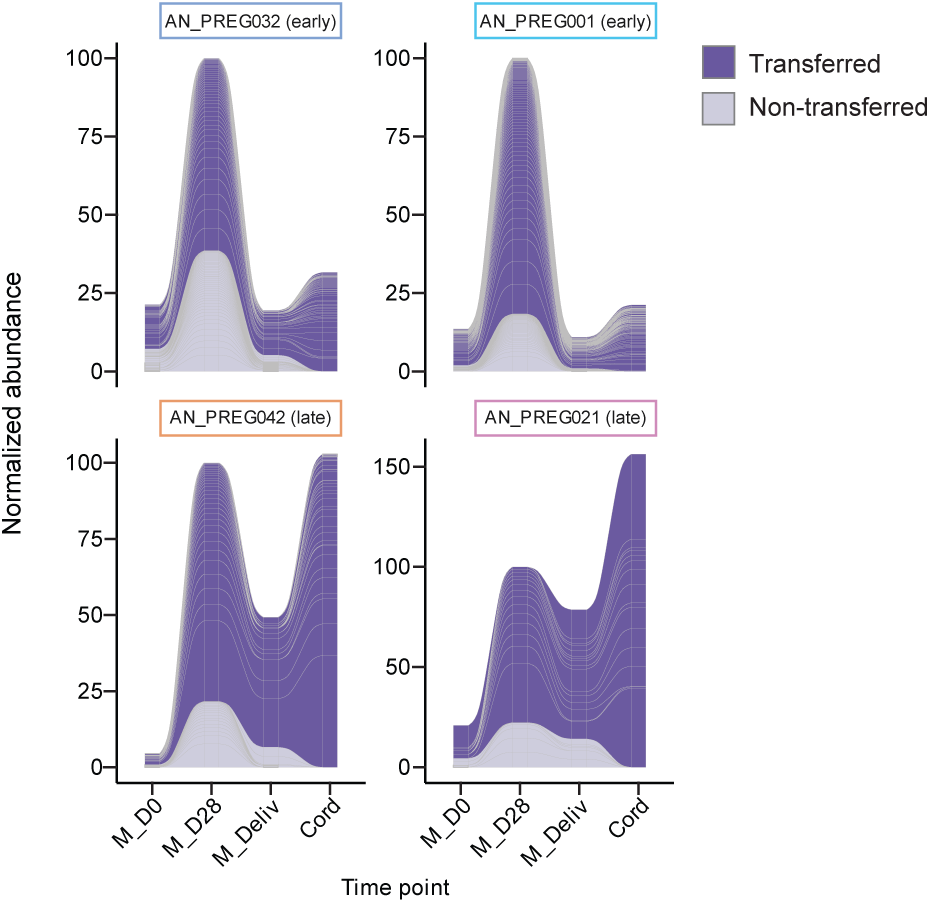
Alluvial plots from four donors. The plots show the abundance of the transferred and non-transferred IgG clonotypes over time, normalized to M_D28.

**Supplementary Figure 5.**
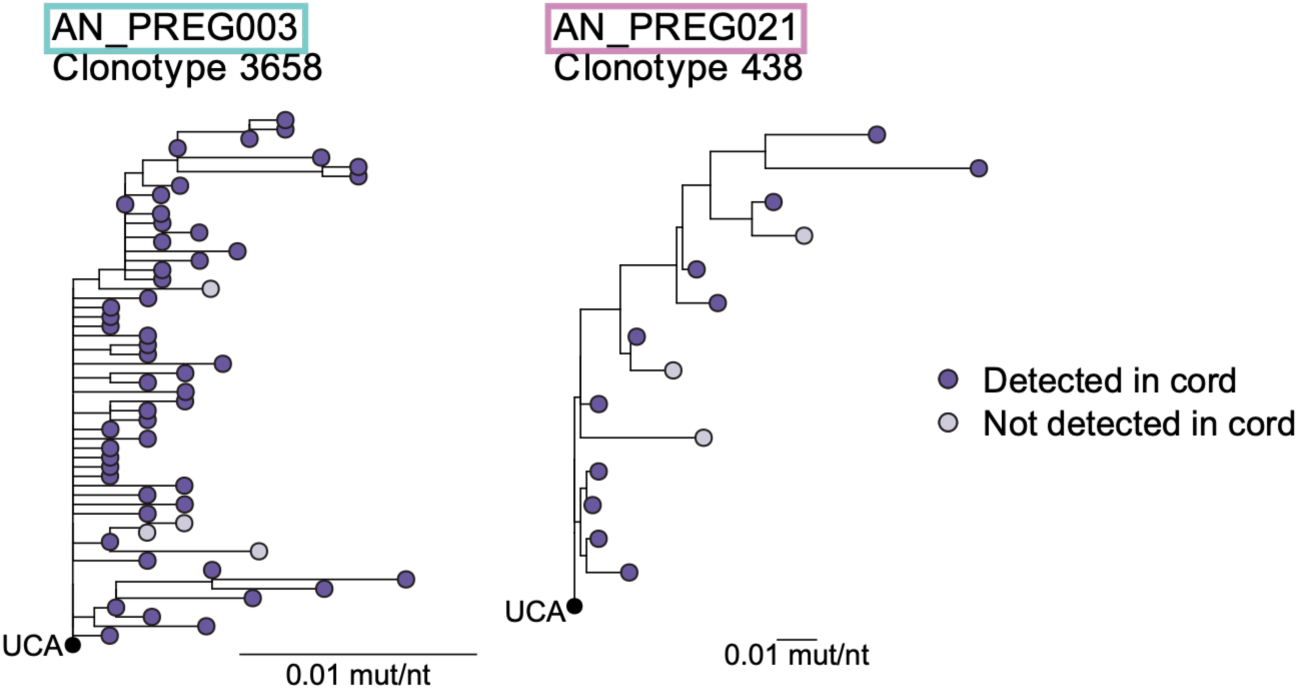
Additional examples of phylogenetic trees of transferred clonotypes. Tip color represents detection in cord. Scale bar represents mutations per nucleotide site. UCA, unmutated common ancestor.

**Supplementary Figure 6.**
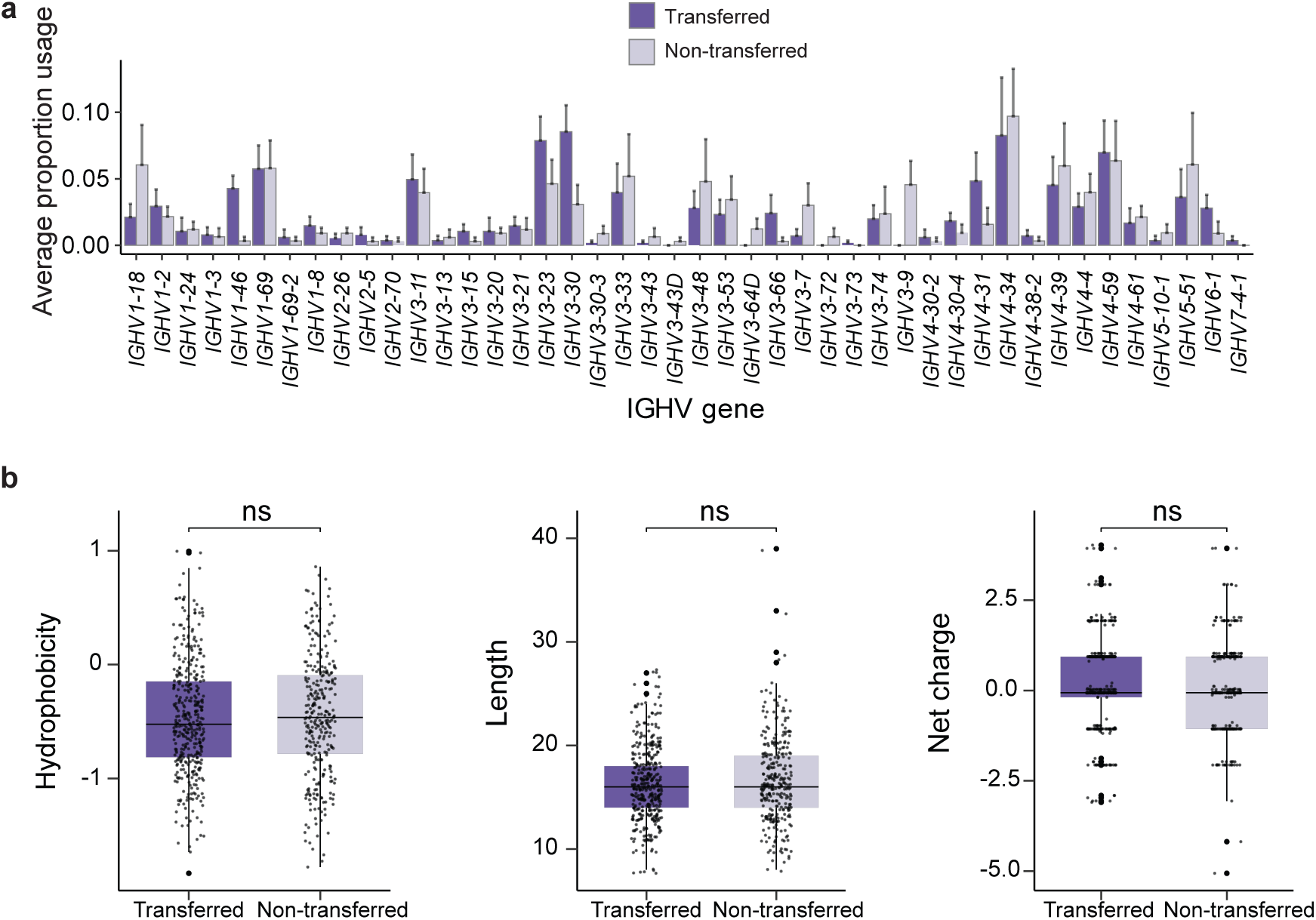
Additional features of the maternal S-specific transferred and non-transferred clonotypes. **a,** *IGHV* gene usage frequency among transferred and non-transferred IgG. Values are mean across six donors; error bars indicate s.e.m. No pairwise comparisons reached significance. **b,** CDRH3 biophysical properties (hydrophobicity, net charge, and length) for transferred and non-transferred IgG clonotypes. Boxplots show median and interquartile range. ns = not significant (Wilcoxon signed-rank test).

**Supplementary Figure 7.**
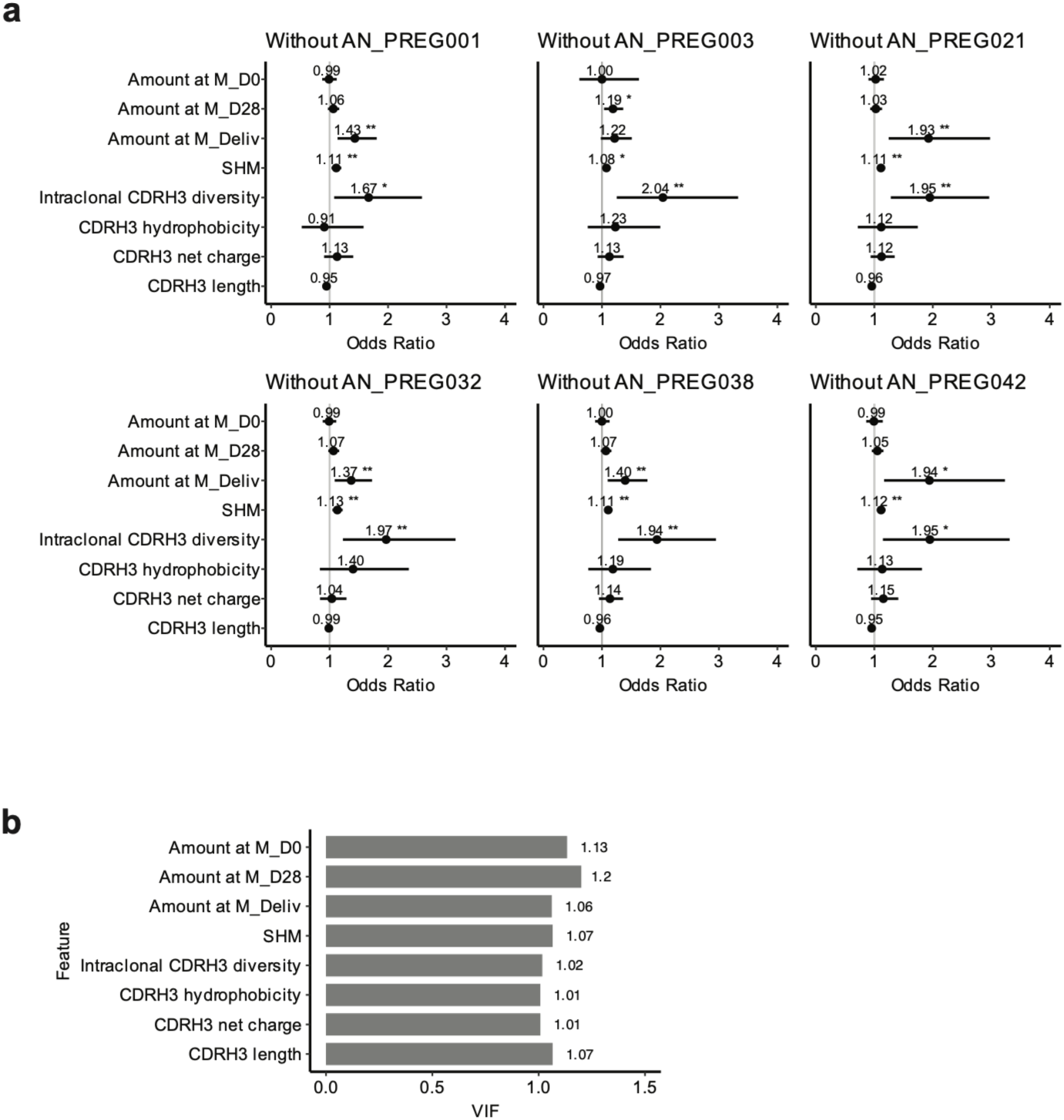
GLMM validation. **a,** GLMM leave-one-out: GLMM modeling effect of predictors on transfer to infant cord for model validation. Each panel represents GLMM run with one donor removed. Consistent results demonstrate that overall effect is not driven by an individual donor. **b,** Variance Inflation Factor (VIF) for model validation. VIF=1 indicates no multicollinearity between predictor variable and all other predictors.

**Supplementary Figure 8.**
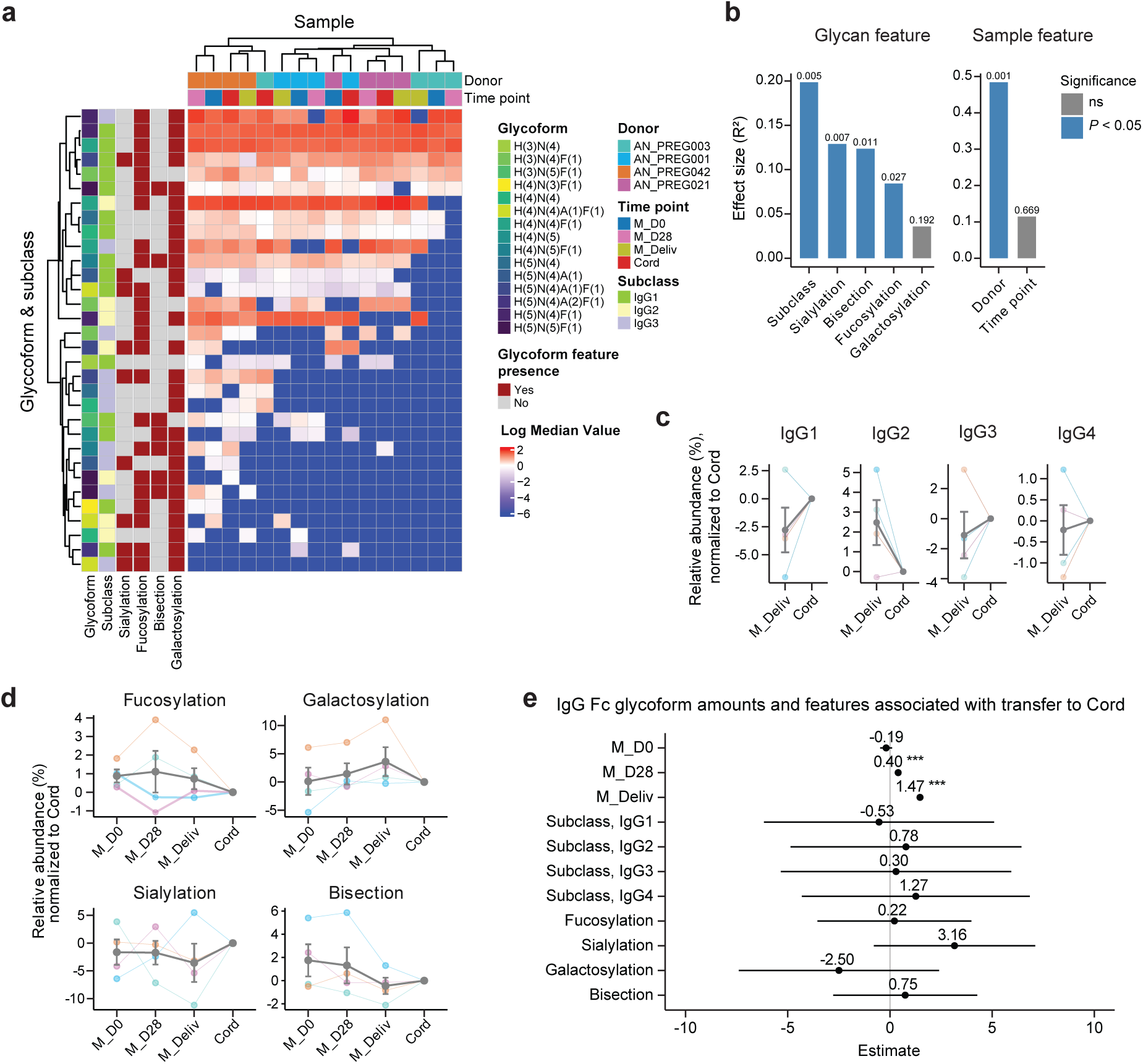
Subclass and Fc glycoform features of S-reactive maternal IgG. **a,** Heatmap of subclass-resolved Fc glycoform (rows) abundance for S-specific IgG across maternal and cord blood samples from four donor dyads (columns). Dendrogram represents results from unsupervised hierarchical clustering. Left annotations denote glycosylation features and top annotations denote donor, time point and IgG subclass. Color scale indicates median-centered log abundance. **b,** The effect size is the PERMANOVA result obtained using the relative abundance values for individual glycoforms. **c,** Relative subclass abundances in maternal-cord dyads at delivery for S-reactive IgG. Thick line represents mean of four donors. Error bars represent s.e.m. **d,** Longitudinal dynamics of S-reactive IgG1 Fc glycosylation features relative to paired cord blood. Thick line represents mean of four donors. Error bars represent s.e.m. **e**, Fixed-effect estimates from a linear mixed-effects model of maternal serum IgG amount (abundance scaled by titer), subclass, and glycosylation features to amount in cord. Vertical line represents null value. Horizontal bars represent 95% CI. Asterisks indicate significance (\*\*\**P* < 0.001).

**Supplementary Figure 9.**
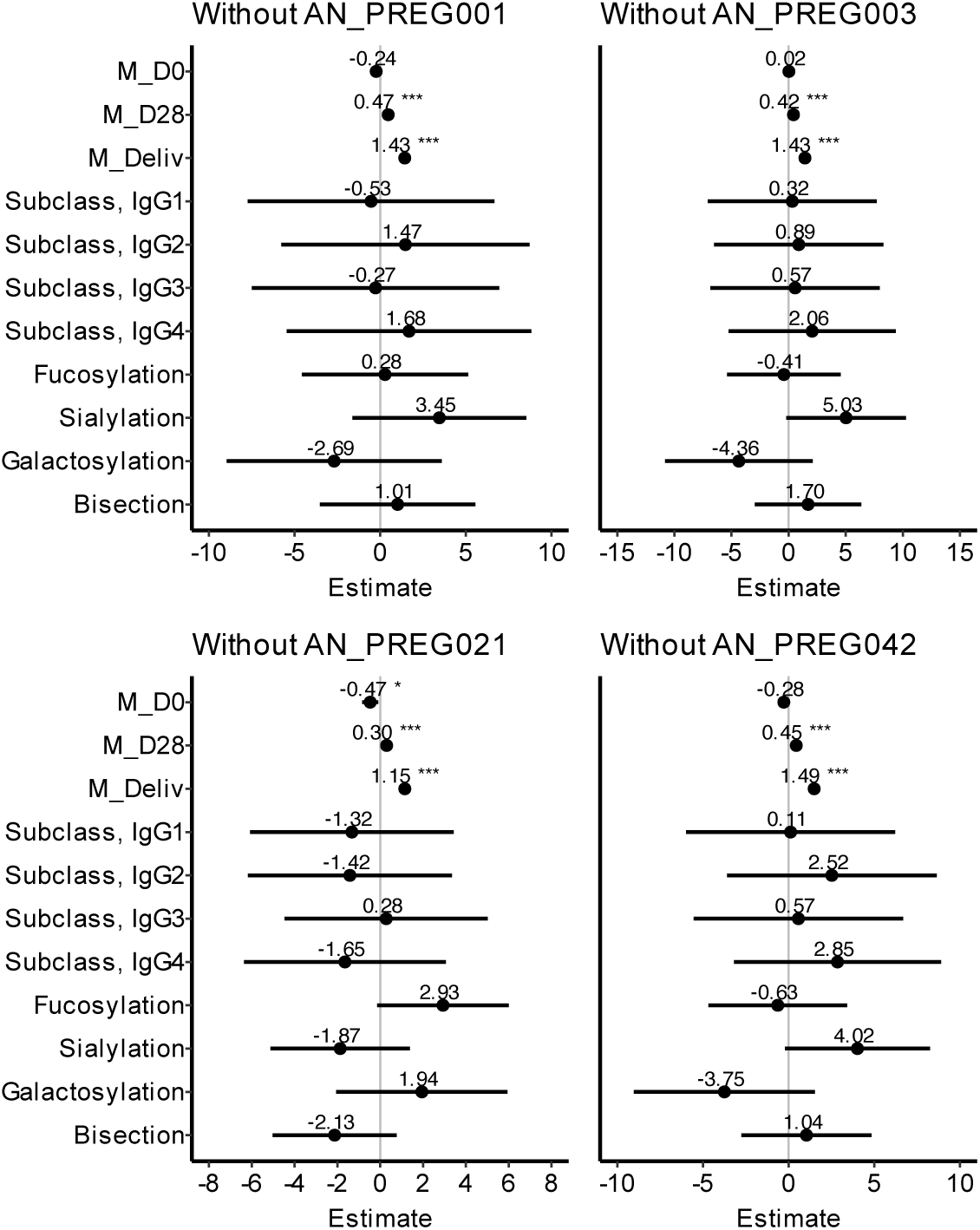
LMM model validation. LMM Minus 1: LMM modeling effect of predictors on transfer to infant cord for model validation. Each panel represents LMM run with one donor removed. Consistent results demonstrate overall effect that is not driven by an individual donor. Asterisks indicate significance (\**P* < 0.05, \*\*\**P* < 0.001).

**Supplementary Figure 10.**
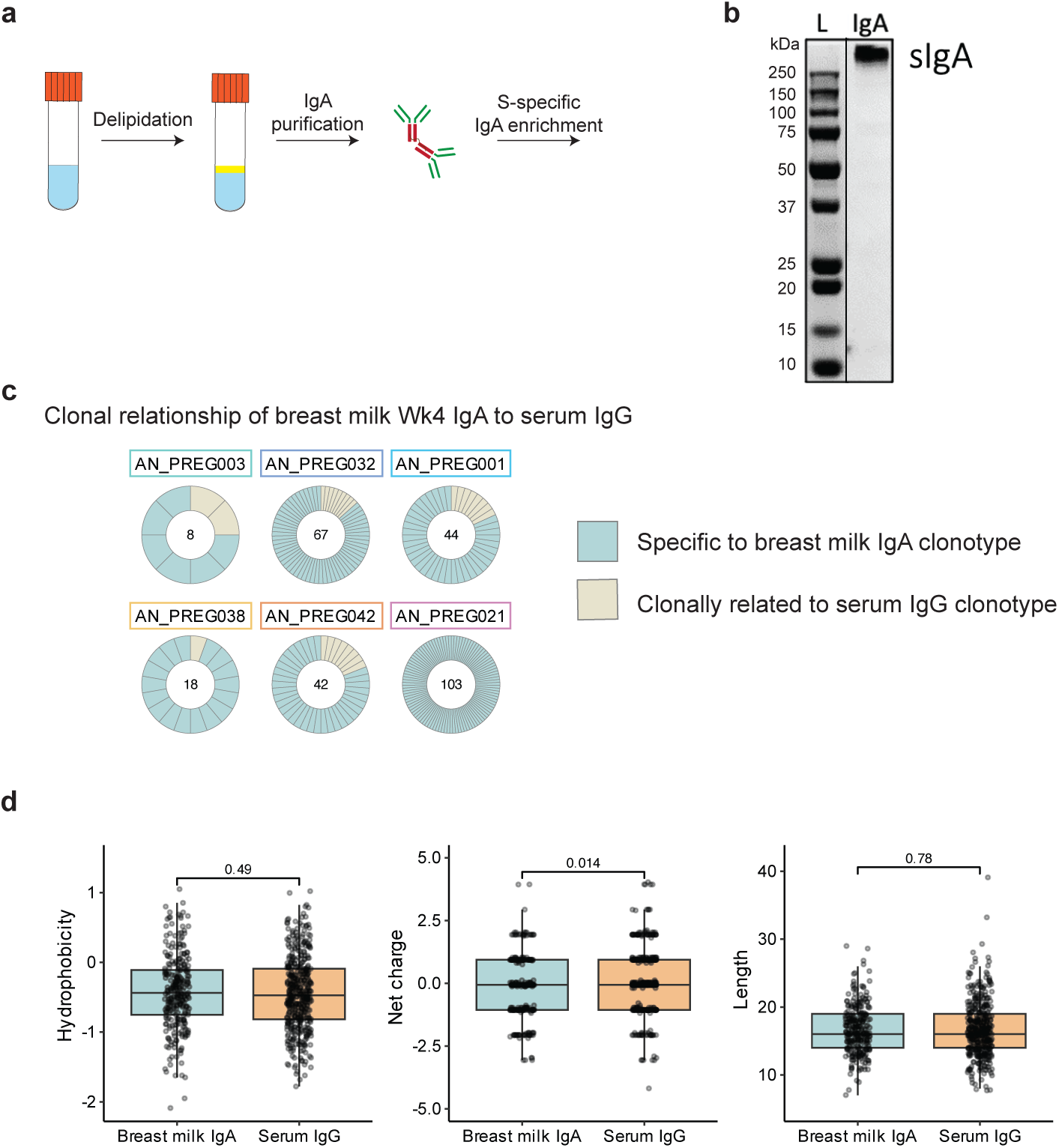
Breast milk IgA purification, validation, and clonotype characterization. **a,** Experimental workflow. Breast milk was delipidated, IgA was purified using Peptide M resin, and S-specific IgA was isolated by affinity pulldown. Full-length IgA was processed for LC-MS/MS-based clonotype identification. **b,** Coomassie-stained SDS-PAGE gel showing a high molecular-weight band for secretory IgA (sIgA). **c,** Donut plots summarizing donor-specific breast milk IgA repertoires by frequency. Each segment represents an individual breast milk IgA clonotype, with clonotypes evenly spaced. Numbers in the center indicate the total number of clonotypes detected. **d,** Biophysical CDRH3 features of serological IgG and breast milk IgA clonotypes. Boxplots show median and quartiles.

